# The phosphorylation landscape of infection-related development by the rice blast fungus

**DOI:** 10.1101/2023.08.19.553964

**Authors:** Neftaly Cruz-Mireles, Miriam Osés-Ruiz, Paul Derbyshire, Clara Jégousse, Lauren S. Ryder, Mark Jave Bautista, Alice Bisola Eseola, Jan Sklenar, Bozeng Tang, Xia Yan, Weibin Ma, Kim C. Findlay, Vincent Were, Dan MacLean, Nicholas J. Talbot, Frank L.H. Menke

## Abstract

Many of the world’s most devastating crop diseases are caused by fungal pathogens which elaborate specialized infection structures to invade plant tissue. Here we present a quantitative mass spectrometry-based phosphoproteomic analysis of infection-related development by the rice blast fungus *Magnaporthe oryzae*, which threatens global food security. We mapped 8,005 phosphosites on 2,062 fungal proteins, revealing major re-wiring of phosphorylation-based signaling cascades during fungal infection. Comparing phosphosite conservation across 41 fungal species reveals phosphorylation signatures specifically associated with biotrophic and hemibiotrophic fungal infection. We then used parallel reaction monitoring to identify phosphoproteins directly regulated by the Pmk1 MAP kinase that controls plant infection by *M. oryzae*. We define 33 substrates of Pmk1 and show that Pmk1-dependent phosphorylation of a newly identified regulator, Vts1, is required for rice blast disease. Defining the phosphorylation landscape of infection therefore identifies potential therapeutic interventions for control of plant diseases.

## INTRODUCTION

Fungal pathogens have evolved specialized infection-related structures to penetrate the tough outer layers of plants to cause disease. These infection structures– which include appressoria, hyphopodia and infection cushions –are important determinants of some of the most serious crop diseases, including cereal rusts and powdery mildews ^1,2^ but their biology is poorly understood. Formation of these structures requires major morphogenetic changes and remodeling of signaling transduction events, including phosphorylation. Some of these events are controlled by major regulators, such as mitogen-activated protein kinases (MAPK). The devastating rice blast fungus *Magnaporthe oryzae*, which destroys enough rice each year to feed 60 million people, develops a melanin-pigmented appressorium that generates enormous turgor pressure of up to 8.0 MPa, which enables the pathogen to rupture the tough rice leaf cuticle ^3,4^. Appressorium morphogenesis in *M. oryzae* requires a MAPK signaling pathway in which the Pmk1 MAPK is a central component ^5^. The importance of Pmk1 is illustrated by the fact that *τ1pmk1* mutants are unable to form appressoria and fail to cause blast disease, while conditional inactivation of the kinase using an analogue-sensitive mutant, prevents invasive fungal growth in plant tissue ^5,6^. Global transcriptional profiling has revealed that Pmk1 regulates 49% of the *M. oryzae* transcriptome during appressorium development– highlighting both its importance, and the complexity of infection-related morphogenesis ^7^.

Components of the Pmk1 cascade have predominantly been identified based on their counterparts in the well-known yeast Fus3/Kss1 pathway required for pheromone signaling and invasive growth ^8,9^. Upstream kinases Mst11 and Mst7, for example, were functionally characterized based on their homology to the yeast MAPKKK Ste11 and MAPKK Ste7, respectively ^10^. Similarly, the adaptor protein Mst50 was identified by homology to Ste50 and shown to control the activity of the three-tiered Mst11-Mst7-Pmk1 MAP kinase module during appressorium formation ^11^. A limited number of downstream interactors of Pmk1 have also been identified, including transcription factors Mst12, Hox7 and Slf1 ^7,12^, and the Pmk1- interacting clone Pic5 ^13^. However, the molecular mechanisms through which these downstream Pmk1 signaling components regulate blast infection remain unknown.

Importantly, Pmk1 counterparts have now been identified in more than 30 fungal pathogen species, including major human, animal, and plant pathogens. In all cases so far reported, these MAPKs have been shown to be necessary for fungal pathogenicity ^14,15^. This includes the causal agents of many of the world’s most significant crop diseases, including Septoria blotch of wheat, southern corn leaf blight, and Fusarium head blight ^16^, and encompasses fungal pathogens exhibiting biotrophic, hemibiotrophic and necrotrophic interactions. Therefore, a common feature of very diverse fungal pathogens, irrespective of whether they cause invasion of living plant tissue or destructive activity to kill plant cells, is their dependence on Pmk1- related MAPK pathways to regulate invasive growth. Collectively, these studies suggest that the Pmk1 MAPK signaling pathway may be a conserved pathway associated with fungal invasive growth which has diversified among distinct groups of pathogens. However, there is little direct evidence for this proposition because the substrates of Pmk1-related MAPKs are largely unknown in any fungal pathogen studied to date.

In this study we decided to take advantage of recent advances in quantitative mass spectrometry (MS) to analyze the global pattern of phosphorylation ^17^ during infection-related development of *M. oryzae*. We set out to define the phosphorylation signature of MAPK signaling ^18^ associated with plant infection by fungal pathogens and identify the cellular signaling pathways regulated by Pmk1 ^19^. Here we report the phosphorylation landscape of appressorium morphogenesis by the blast fungus and define the major changes in phosphorylation that occur during fungal development. We use this resource to identify conserved phosphosites specific to fungal pathogens that elaborate diverse infection structures and which exhibit distinct modes of fungal pathogenesis– including biotrophic, hemibiotrophic and necrotrophic species – thereby defining the putative patterns of MAPK signaling across 40 major disease-causing fungal species. To validate this approach, we identified 201 phosphosites and classified them into signaling pathways and physiological processes required for infection by the blast fungus and used parallel reaction monitoring to identify 33 novel putative Pmk1 substrates. This analysis enabled the identification of a new regulator of appressorium morphogenesis, Vts1, which requires Pmk1-dependent phosphorylation to fulfil a key role in rice blast disease. When considered together, this study provides the most comprehensive analysis of infection- phosphorylation by a fungal pathogen to date and highlights how phosphoproteomic analysis can provide unprecedented insight into the biology of fungal invasive growth.

## RESULTS

### The Pmk1 MAP kinase is activated during infection-related development by *M. oryzae*

We first set out to define a time-course for global phosphoproteomic analysis of infection-related development by the blast fungus, by identifying the precise time of Pmk1 MAPK activation. Plant infection by *M. oryzae* is initiated when a fungal spore, called a conidium, lands on a hydrophobic surface and germinates to produce a polarized germ tube within 2h (**Figure 1A**). By 4h an incipient appressorium is formed and the contents of the conidium are recycled by autophagy to allow development of the appressorium (**Figure 1A and 1B**). The Pmk1 MAPK is essential for both appressorium development and virulence ^5^, and Δ*pmk1* mutants are unable to form appressoria and cannot infect rice plants (**Figure 1B and 1C**). To investigate the temporal dynamics of Pmk1 activation we germinated spores on an artificial hydrophobic surface that mimics the host leaf surface. We extracted protein from synchronized infection structures from the wild-type *M. oryzae* strain Guy11 and an isogenic Δ*pmk1* mutant. We observed that Pmk1 is phosphorylated on its TEY motif within 1h of conidial germination on a hydrophobic surface (**Figure 1A and 1D**) and remains active for up to 4h. Pmk1 activation therefore precedes infection-related development but is maintained throughout appressorium morphogenesis.

**Figure 1.**
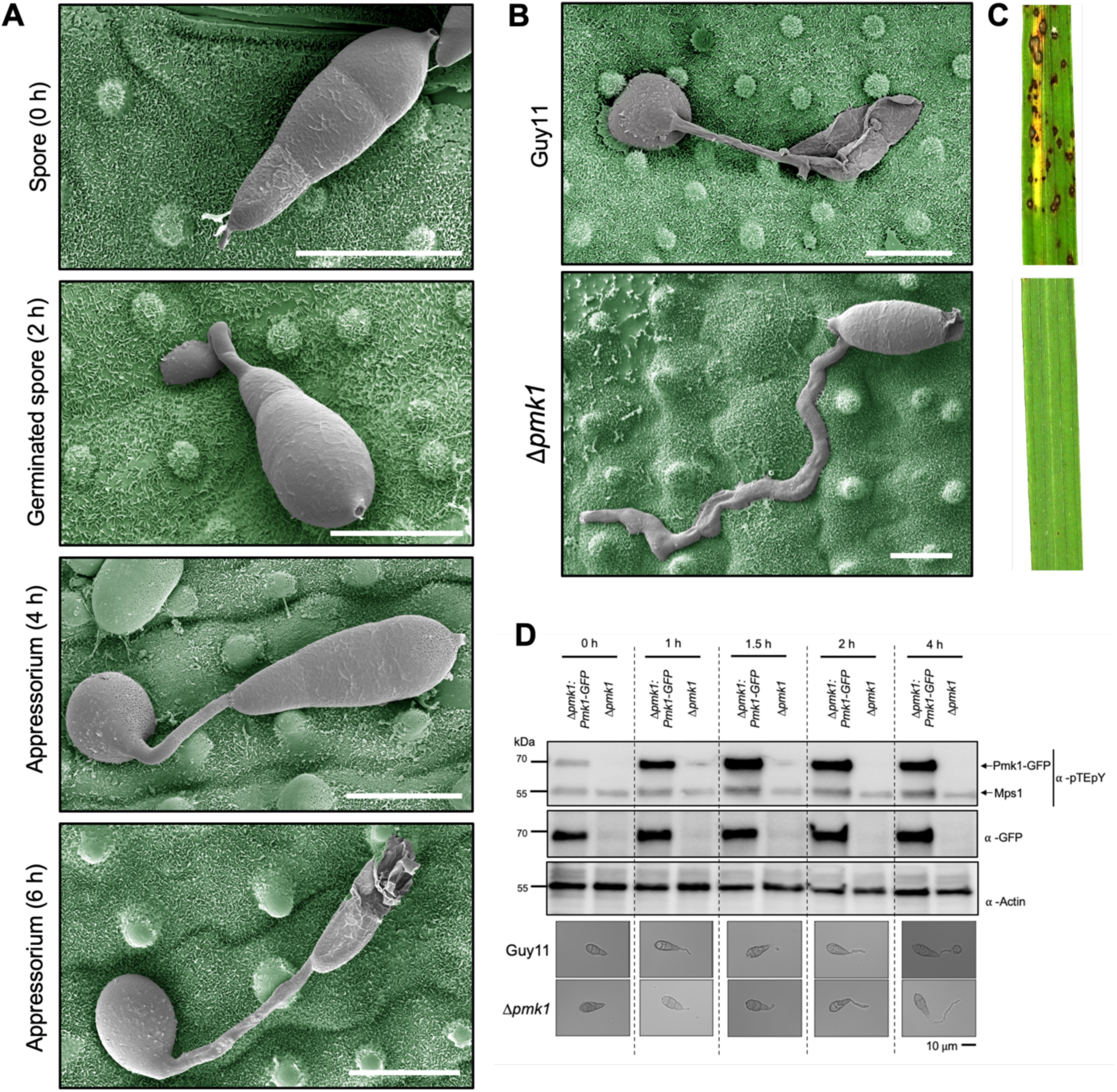
Early morphogenetic transitions in the appressorium overlap with activation of the Pmk1 signaling cascade. **(A)** Scanning electron micrographs (SEM) with false colouring to show appressorium germination of wild-type strain Guy11 at 0h, 2h, 4h and 6h. **(B)** SEM with false colouring to show appressorium germination of Guy11 and 11*pmk1* strains at 24h. In **(A)** and **(B)** the blast fungus is shown in grey, and rice leaf surface is false-colour imaged to green. Scale bars indicate 10 μm. **(C)** Rice leaves infected with Guy11 (top) and 11*pmk1* (bottom) strains. Rice seedlings of cultivar CO-39 were spray-inoculated with conidial suspensions of equal concentrations of each strain and incubated for 5d. **(D)** Western blot analysis of total protein extracted from *in vitro* germinated spores at 0, 1, 1.5, 2 and 4 h from 11*pmk1* complemented with *PMK1-GFP* and 11*pmk1* using ⍺-pTEpY (top panel), ⍺-GFP (middle panel) and ⍺-Actin (lower panel). ⍺- pTEpY has been also reported to detect the MAPK Mps1 ^49^. Proteins were immunoblotted with appropriate antisera (listed on the right). Arrows indicate expected band sizes.

### Infection related development coincides with large changes in the phosphoproteome

To define the phosphorylation landscape of infection-related development by *M. oryzae,* we incubated conidia of Guy11 and the Δ*pmk1* mutant on a hydrophobic surface and performed a large-scale quantitative phosphoproteomics experiment, extracting phosphoproteins from synchronized infection structures at 0, 1, 1.5, 2, 4 and 6h post-germination (**Figure 2A and S1**). Using discovery proteomics based on data-dependent acquisition (DDA), we identified 8005 phosphopeptides from 2,062 proteins during the 6h time-course in both strains (**Figure 2A and 2B**). We quantified this dataset using a label-free MS1-quantification approach (LFQ) and were able to quantify 7048 phosphopeptides. To identify differential phosphopeptides we determined the ratio between Guy11 timepoints compared to conidia (t=0) and filtered for 2- fold change and an adjusted p-value =< 0.05. With these settings, the abundance levels of 5058 phosphopeptides were found to be significantly different during germling (1h, 1.5h, 2h) and appressoria (4h and 6h) stages compared to conidia (0h) in Guy11. As early as 1h after germination, we identified large changes in abundance of phosphopeptides (420 less abundant and 2049 more abundant) and phosphoproteins (100 less abundant and 497 more abundant) (**Table 2**). The phosphorylation landscape of the emerging germling therefore undergoes significant changes due to a combination of changes in the amount of each protein as well as differential phosphorylation. Consistent with our immunoblot analysis of Pmk1 phosphorylation (**Figure 1D**), we can detect phosphorylation of the TEY motif in the activation loop of Pmk1 as early as 1h after germination, peaking at 1.5h and remaining at a sustained level up to 4h, and this phosphorylation was absent in the 11*pmk1* mutant, as expected (**Figure 2C**). At 6h, TEY phosphorylation increases again, suggesting a second Pmk1 activation event occurs between 4-6h, when the appressorium develops and a significant change in growth polarity occurs. As expected, in the 11*pmk1* mutant we cannot detect phosphorylation of the TEY motif at any timepoint. During the first 2h the number of upregulated phosphopeptides in

**Figure 2.**
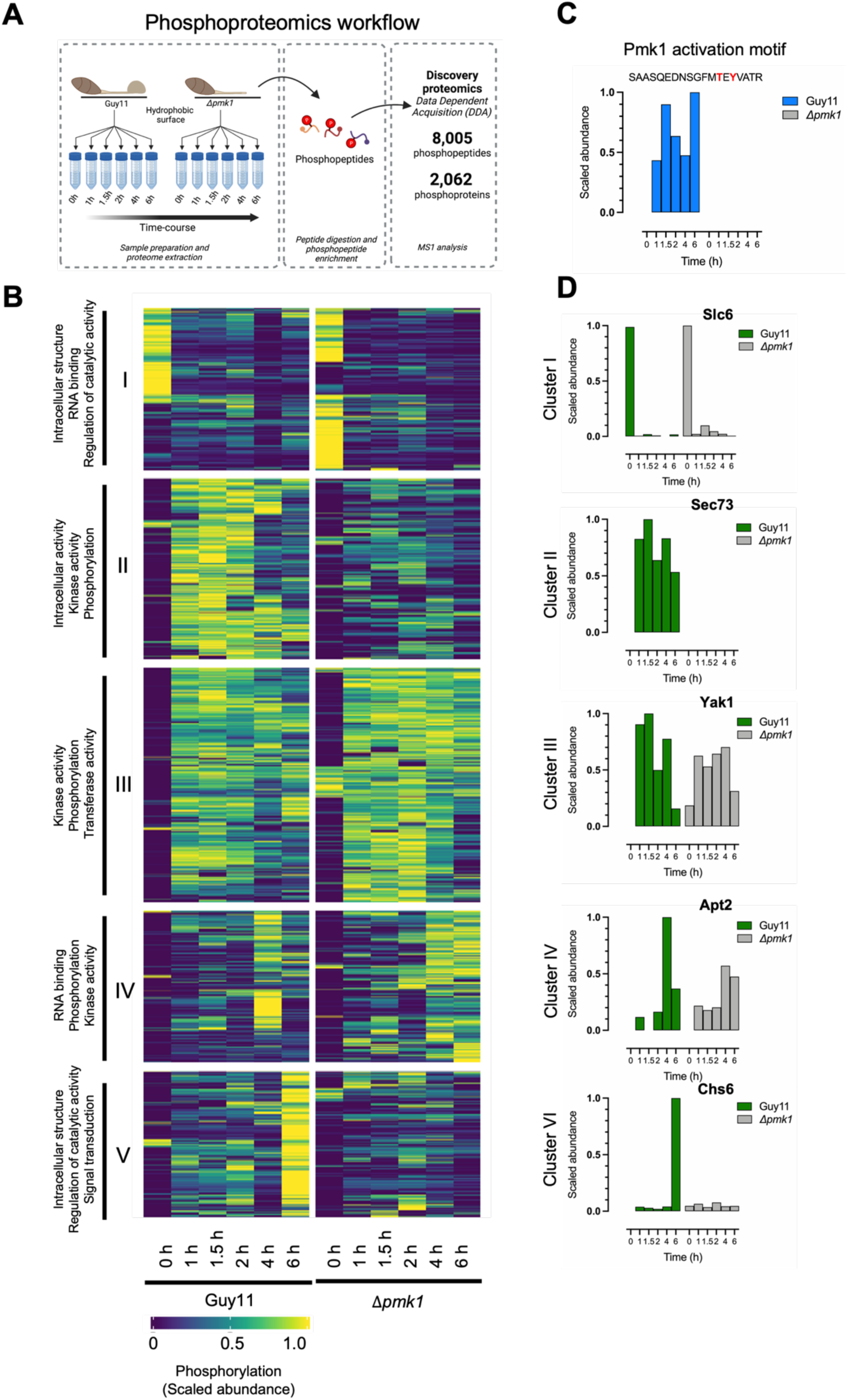
Infection-related development causes major changes in protein phosphorylation. **(A)** Schematic to show the infection-related development experiment in a pathogenic (Guy11) and non-pathogenic (11*pmk1*) *M. oryzae* strain. Spores and germinated cells were synchronized at 0h, 1h, 1.5h, 2h, 4h and 6h. Phosphopeptides were enriched using TiO2 microspheres and were subjected to liquid chromatography-tandem mass spectrometry (LC-MS/MS). Samples were analyzed on Data Dependent Acquisition (DDA) mode to be quantified using an label free quantification based on precursor ions (MS1). **(B)** Heatmap to show the differentially phosphorylated phosphopeptides in Guy11 and 11*pmk1.* Each row corresponds to a single phosphosite and rows are ordered by hierarchical clustering (I-V). The top three GO terms for each cluster are indicated. **(C)** Bar graphs to show relative phosphorylation abundance for the peptide containing the Pmk1 activation motif (pTEpY) during early infection. **(D)** Relative phosphorylation for representative phosphopeptides in each defined cluster. Cluster I, solute carrier family protein Slc6 (MGG_05433); Cluster II, Sec73 (MGG_06905); Cluster III, Yak1 (MGG_06399); Cluster IV, Apt2 (MGG_07012) and Cluster V, chitin synthase 6, CH6 (MGG_13013).

**Table 2.** The number of phosphosites at each developmental stage clustered as defined in. **Figure 2B**.

| Time | Guy11 | | $\Delta pmk1$ | |
| --- | --- | --- | --- | --- |
|  | Up-regulated | Down-regulated | Up-regulated | Down-regulated |
| 1h | 420 | 2049 | 727 | 1681 |
| 1.5h | 469 | 2168 | 619 | 1844 |
| 2h | 344 | 2332 | 527 | 1986 |
| 4h | 599 | 2323 | 569 | 2373 |
| 6h | 329 | 3322 | 744 | 1736 |

11*pmk1* is significantly lower (1681), compared to the wild type, whilst the number of downregulated phosphopeptides is significantly higher (727) consistent with the absence of Pmk1 activity (**Figure 2C**). K-means clustering of the data revealed 5 clusters enriched for differential phosphopeptides in conidia (Cluster I), germinated conidia (Cluster II and Cluster III) and incipient appressoria (Cluster IV and Cluster V) (**Figure 2B**). Changes in overall phosphopeptide abundance are similar for Guy11 and the 11*pmk1* mutant in clusters I, III and IV, while the abundance changes in clusters II and V are drastically different between Guy11 and 11*pmk1*. The absence or significant reduction in phosphopeptide abundance in the 11*pmk1* background is consistent with high Pmk1 activity in Guy11 at those times. GO term enrichment analysis for 4 of these clusters revealed biological processes and molecular functions related to signal transduction and protein phosphorylation (**Figure S2**). Whilst cluster I is enriched for intracellular anatomical structures, RNA binding, proteins associated with plasma membrane and cation transport, but not signal transduction or protein phosphorylation. This indicates that protein phosphorylation is one of the key primary processes during infection-related development. GO terms related to other processes known to be essential for appressorium development in *M. oryzae* are also enriched, such as autophagy (cluster II), lipid binding and actin binding (cluster III) and microtubule and intracellular transport (cluster IV) ^20–23^. Cluster I represents phosphopeptides derived from proteins that are present in conidia and show lower relative abundance during the course of infection-related development. These include proteins such as the putative solute carrier transporter Slc6 (MGG_05433) (**Figure 2D**), Glycerate-3- kinase (MGG_06149), Linolate diol synthase LDS1 (MGG_10859) and eukaryotic translation initiation factor 3 subunit a (MGG_10192) (**Figure S2**), most of which have been shown to have a function in virulence in *M. oryzae* ^24,25^, but which represent proteins involved in metabolic functions rather than signaling. Cluster III represents proteins phosphorylated early during appressorium development with high relative abundance of the corresponding phosphopeptides between 1-4h in both Guy11 and the 11*pmk1* mutant, including the Yak1 kinase (MGG_06399) (**Figure 2D**) and protein phosphatase Ssd1 (MGG_08084), both of which are required for virulence ^26,27^. Phosphopeptides in Cluster II show a similar relative abundance at early timepoints in Guy11 but little to no increase in abundance in the 11*pmk1* mutant, suggesting the corresponding proteins require Pmk1 for their phosphorylation or expression. Examples of phosphopeptides showing this type of pattern include peptides from protein transporter Sec73 (MGG_06905) (**Figure 2D)** and kinases Sch9 (MGG_14773) and Atg1 (MGG_06393) (**Figure S2)**. Cluster IV and V represent phosphopeptides with high relative abundance at the time when appressorium begins to form and these include the aminophospholipid translocase Apt2 (MGG_07012), which is essential for plant infection ^28^, the transcription factor Hox7 (MGG_12865), implicated in appressorium morphogenesis ^7^, chitin synthase Chs6 (MGG_13013) and the glucosamine-6-phosphate isomerase (MGG_00625). The latter two proteins represent cluster V which again shows phosphopeptides with high abundance at 6h in Guy11 but mostly absent from the 11*pmk1* mutant, consistent with Pmk1-dependent phosphorylation or expression. Taken together, these results show that during infection-related development by the blast fungus, major changes in the phosphoproteome landscape are occurring, including many proteins known to be required for virulence, and a subset of these phosphoprotein changes require presence of the Pmk1 kinase.

### Distinct patterns of phosphosite conservation are evident across fungal species showing diverse modes of pathogenesis

Given the extensive changes in phosphorylation during infection-related development by *M. oryzae*, we decided to investigate the extent of phosphosite conservation across different fungal species. Using Orthofinder we identified orthogroups for 41 filamentous fungal species including saprophytes, mutualists, plant pathogens, and human pathogens, as well as *Saccharomyces cerevisiae* and *Schizosaccharomyces pombe*, the two model yeast species. We then mapped the conservation of all our identified *M. oryzae* phosphorylated residues onto the orthogroups for all 41 fungal species, *k*-means clustered them, using the elbow method according to the subset of species in which residues were conserved, and visualized the data in a heatmap containing 9 *<u>C</u>onserved <u>P</u>hosphorylated <u>R</u>esidue* (CPR) groups (**Figure 3**) . We identified a total of 1,198 conserved phosphorylated residues. Clustering species based on the conservation of phosphorylated residues (**Figure 3A**) results in a tree composed of distinct clades, when compared to a phylogenetic tree based on the orthogroups (**Figure 3C**). This provides evidence that a large proportion of the conservation of phosphorylated residues observed cannot be explained by protein conservation at the amino acid level or phylogenetic distance between species. The heatmap shows several interesting CPR clusters of which Cluster 4 likely represents core signaling proteins and conserved phosphorylated residues observed in the majority of fungal species (**Figure 3B**). Consistent with this, the cluster includes 69 proteins of which 15 are annotated as protein kinases, including the MAPKs Pmk1 and Osm1 and the MAPKK Mst7. Interestingly, CPR Cluster 9 shows conservation of phosphorylated residues among the majority of plant pathogenic species but only limited conservation in saprophytes such as *Aspergillus nidulans, A. fumigatus* and *A. flavus*, suggesting that these residues are associated with a lifestyle dependent on plant hosts. As expected, these two clusters show a strong correlation with a requirement for a Pmk1 MAPK orthologue for virulence/pathogenicity (**Table 1**). CPR Cluster 6 shows a strong correlation with 16 mainly hemibiotrophic plant pathogenic fungal species– hemibiotrophic species are those that invade living plant tissue initially in their life cycle, before killing plant cells at later stages of development. This includes many of the most important crop disease-causing fungi, such as *M. oryzae* and *Zymoseptoria tritici* and reveals phosphosite conservation in at least 13 transcription factors and a wide range of metabolic enzymes (**Table S1**) Cluster 3 stands out because it appears tightly associated with the genus *Fusarium*, including the wheat head blight pathogen *F. graminearum* for example, and shows phosphosite conservation in pH-responsive and morophogenetic transcriptional regulators, for example. Cluster 5 shows high levels of conservation among Dothideomycete pathogens, including the causal agent of new blotch of barley *Pyrenophora teres,* brown spot of rice, *Bipolaris oryzae*, and southern corn leaf blight, *Cochliobolus heterostrophus*. The conservation of phosphosites within this group of related cereal pathogens, provides evidence that the regulation of invasive growth may be similarly configured to *M. oryzae.* Finally, Cluster 2 shows phosphosites conserved solely among fungi producing pressurized, melanin-pigmented appressoria, such as the *Colletotrichum* anthracnose pathogens, and consistent with this shows phosphosite conservation in a range of proteins previously implicated in appressorium morphogenesis (**Table S1**). When considered together, this comparative analysis provides evidence that subsets of the identified phosphorylated residues in *M. oryzae* are conserved among diverse pathogens showing some common life-style features, such as plant-association, invasive growth in living plant tissue, and the formation of specialized infection structures, providing a key resource for defining the signaling mechanisms that govern fungal pathogenesis.

**Figure 3.**
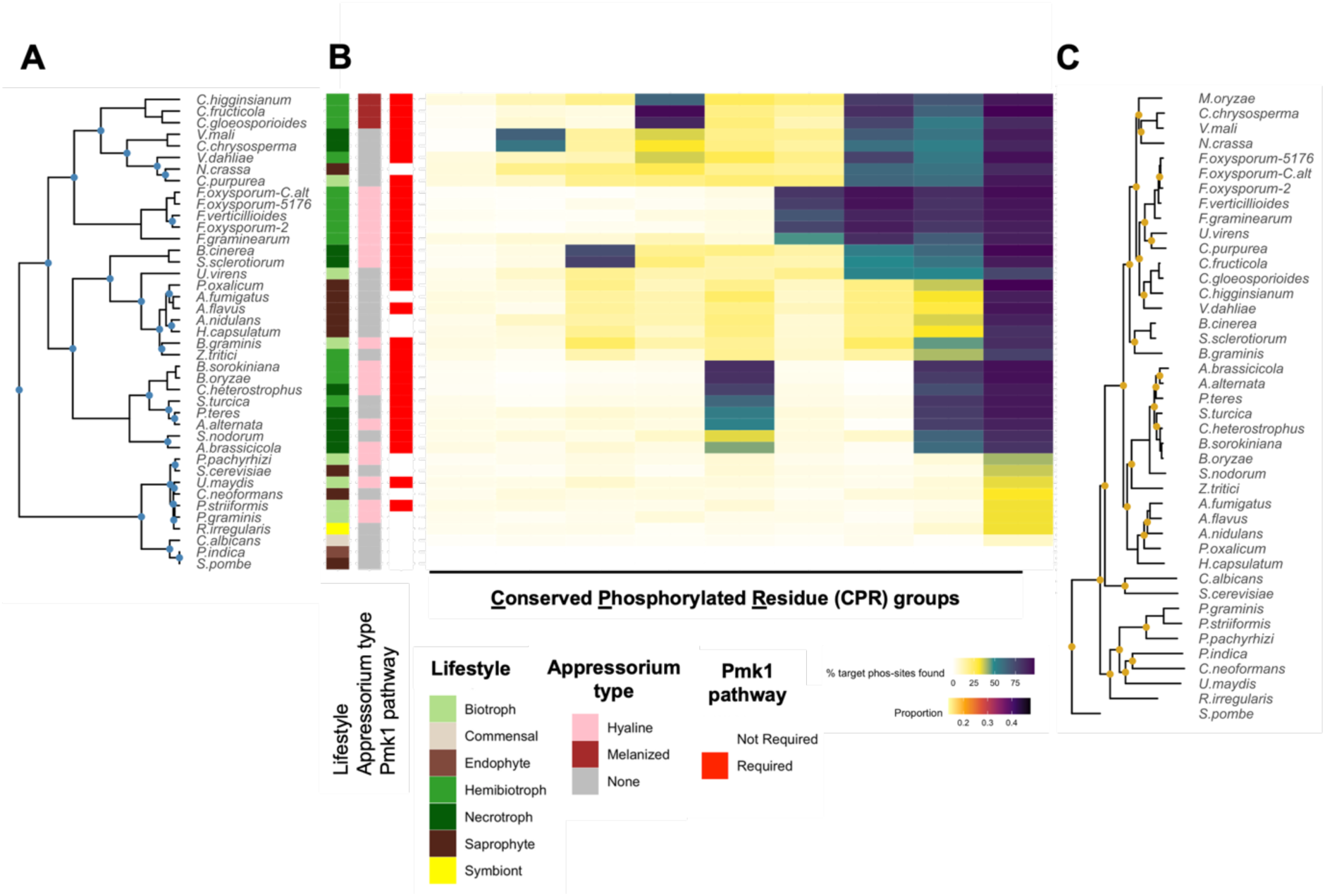
Evolutionary analysis determines appressorium specific phosphosites across fungal species. **(A)** Fungal species tree based on based on the conservation of phosphorylated residues. **(B)** Heatmap to show *k*-means clustered conserved phosphorylated residues among 41 fungal species from the infection-related development dataset in *M. oryzae*. Each row corresponds to a single species, and rows and columns are ordered by hierarchical clustering in 9 <u>C</u>onserved <u>P</u>hosphorylated <u>R</u>esidue (CPR) groups. Colour indicates the percent of target phosphosites from *M. oryzae* found in other species. **(C)** Phylogenetic tree based on the orthogroups for the 41 fungal species: *Alternaria alternata, Alternaria brassicicola, Aspergillus flavus, Aspergillus fumigatus, Aspergillus nidulans, Bipolaris oryzae, Bipolaris sorokiniana (Cochliobolus sativus), Blumeria graminis, Botrytis cinerea, Candida albicans, Claviceps purpurea, Cochliobolus heterostrophus, Colletotrichum fructicola, Colletotrichum gloeosporioides, Colletotrichum higginsianum, Cryptococcus neoformans, Cytospora chrysosperma, Fusarium graminearum, Fusarium oxysporum-2, Fusarium oxysporum-5176, Fusarium oxysporum- C.alt, Fusarium verticillioides, Histoplasma capsulatum, Neurospora crassa, Penicillium oxalicum, Phakopsora pachyrhizi, Piriformospora indica, Puccinia graminis, Puccinia striiformis, Pyrenophora teres, Rhizophagus irregularis, Saccharomyces cerevisae, Schizosaccharomyces pombe, Sclerotinia sclerotiorum, Setosphaeria turcica, Stagonospora nodorum, Ustilaginoidea virens, Ustilago maydis, Valsa mali, Verticillium dahliae, Zymoseptoria tritici*.

**Table 1.** List of characterized Pmk1 orthologs in pathogenic fungi.

| Fungal species | Fus3/Kss1 orthologue | Reference |
| --- | --- | --- |
| *Alternaria alternata* | Fus3 | 50 |
| *Alternaria brassicicola* | Amk1 | 51 |
| *Aspergillus flavus* | MpkB | 52 |
| *Bipolaris oryzae* | Bmk1 | 53 |
| *Bipolaris sorokiniana*<br>(*Cochliobolus sativus*) | Fus3 | 54 |
| *Blumeria graminis* | Mpk1 | 55 |
| *Botrytis cinerea* | Bmp1 | 56 |
| *Claviceps purpurea* | Cpmk1 | 57 |
| *Cochliobolus heterostrophus* | Chk1 | 58 |
| *Colletotrichum fruticicola* | Cfmk1 | 59,60 |
| *Colletotrichum gloeosporioides* | CgMk1 | 61 |
| *Colletotrichum higginsianum* | ChMK1 | 62 |
| *Colletotrichum lagenarium* | Cmk1 | 63 |
| *Colletotrichum scovillei* | CsPmk1 | 64 |
| *Cytospora chrysosperma* | CcPmk1 | 65 |
| *Fusarium graminearum* | Gmpk1 | 66 |
| *Fusarium oxysporum* | Fmk1 | 67 |
| *Fusarium verticillioides* | Mk1 | 68 |
| *Magnaporthe oryzae* | Pmk1 | 5 |
| *Metarhizium robertsii* | Fus3 | 69 |
| *Mycosphaerella graminicola* | Fus3 | 70 |
| *Penicillium oxalicum* | PoxMk1 | 71 |
| *Puccinia striiformis* | Mapk1 | 72 |
| *Pyrenophora teres* | Ptk1 | 73 |
| *Sclerotinia sclerotiorum* | Smk1 | 74 |
| *Setosphaeria turcica* | Stk2 | 75 |
| *Stagonospora nodorum* | Mak2 | 76 |
| *Ustilaginoidea virens* | UvPmk1 | 77 |
| *Ustilago maydis* | Kpp2 Kpp6 | 78,79 |
| *Valsa mali* | VmPmk1 | 80 |
| *Verticillium dahliae* | Vmk1 | 81 |

### Phosphorylation of signaling pathways controlling infection-related development

To investigate the relevance of the observed patterns of phosphorylation on infection-related development, we mapped 201 differentially phosphorylated residues identified by LFQ onto proteins in signaling pathways implicated in appressorium morphogenesis and plant infection. Out of 17 proteins associated with the Pmk1 MAPK signaling pathway, 11 proteins show abundance changes for 1 or more phosphopeptides, including proteins acting upstream of Pmk1, such as the MAPKK Mst7 and adaptor protein Mst50 ^10,11,29,30^, as well as potential and verified downstream Pmk1 targets, such as transcription factors Sfl1, Znf1, Hox7 and Mst12 ^7,12,31,32^ (**Figure 4A**). This high proportion of infection-regulated phosphoproteins is not limited to the Pmk1 pathway because 12 out of 25 proteins mapped onto the Sln1 histidine kinase signaling pathway, required for appressorium turgor sensing ^3^, show phosphopeptide abundance changes over the 6h time course of infection related development (**Figure 4D and S4**). Autophagy is known to be required for appressorium function and dependent on Pmk1 ^20,33^. Out of 23 proteins involved in autophagy, 8 proteins associated with initiation and selective autophagy show abundance changes on one or more phosphopeptides (**Figure 4E and S4**). In the Cyclic AMP Protein kinase A-dependent signaling pathway, which acts in concert with Pmk1 to regulate initiation of appressorium development and turgor generation, 3 proteins show change in phosphoprotein levels (**Figure 4B and S4**). While the recently described Vast1 pathway, implicated in control of appressorium maturation has 4 of 5 Vast1 phosphoproteins changing in abundance, predicting that this pathway is key to plant infection (**Figure 4C and S4**). In all of the signaling pathways analysed the phosphopeptide abundance profiles are distinct between Guy11 and the 11*pmk1*mutant, providing evidence that Pmk1 plays a role in their direct or indirect regulation, and highlighting the global nature of its regulatory effect on physiological and morphogenetic processes necessary for elaboration of a functional appressorium.

**Figure 4.**
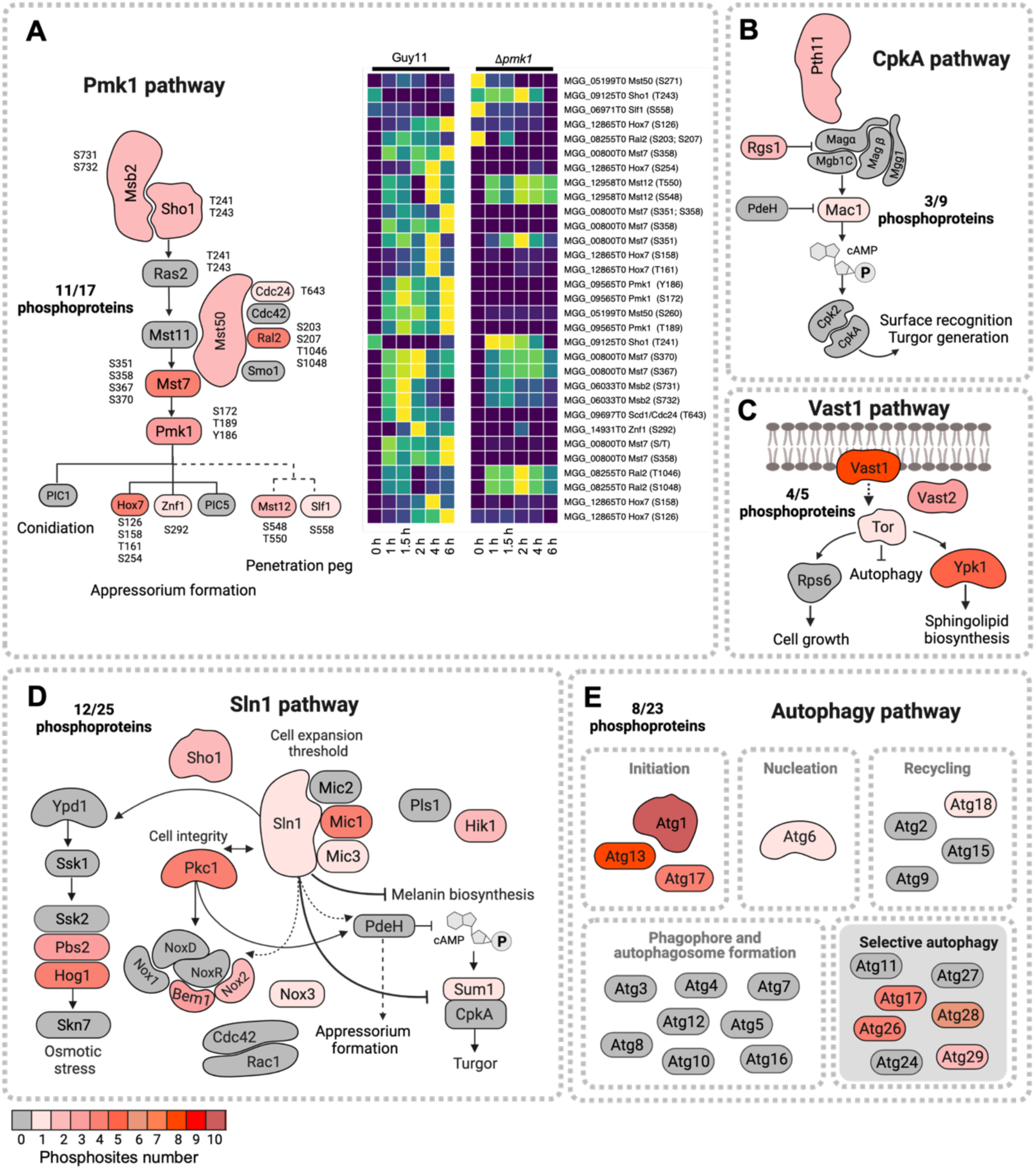
Phosphorylation is a major modification in proteins from pathways that control infection-related development. **(A)** On the left, schematic representation of components of the Pmk1 signaling pathway in *M. oryzae*. Differentially phosphorylated residues in are indicated for each protein (p value <= 0.05). On the right, heatmap to show the differentially phosphorylated phosphopeptides in Guy11 and 11*pmk1* during infection-related development. Each row corresponds to a single phosphosite. **(B-E)** Schematic representation of components of (B) CpkA, (C) Vast1, (D) Sln1 and (E) Autophagy pathways. For all represented proteins, the number of differentially phosphorylated residues is represented in red.

### Targeted phosphoproteomics defines potential targets of the Pmk1 MAPK

Given the importance of Pmk1 in regulation of proteins required for plant infection by *M. oryzae*, we decided to identify direct substrates of the MAPK. For this purpose, we carried out targeted quantitative phosphoproteomic analysis in *M. oryzae* Guy11 and the 11*pmk1* mutant by Parallel Reaction Monitoring (PRM). We used this separate, more accurate approach to enhance confidence in the identification of differential abundance of phosphopeptides as well to benchmark the LFQ data (**Supplemental Data 1**). We hypothesized that direct targets of Pmk1 would be phosphorylated only in the presence of Pmk1. For PRM, we selected peptides to target from a DDA library using the following criteria: a) differentially phosphorylated peptides based on LFQ; b) peptides from previously reported Pmk1 target proteins; and c) potential components of the Pmk1 pathway based on a hierarchical transcriptomic analysis ^7^. Using this information, we selected a list of 286 phosphopeptides belonging to 101 proteins (**Figure 5A**). Of the 286 phosphopeptides quantified by PRM, 182 showed differential abundance compared to conidia in one or both genotypes, representing 86 proteins (fold change: -1< log2(FC) >1; p-value =<0.05) (**Figure S5)** . Of the 182 phosphopeptides showing differential abundance, 63 peptides from 33 proteins were differentially phosphorylated at one or more timepoints in Guy11 and non-differentially phosphorylated in the 11*pmk1* mutant (**Figure 5B**). We named these proteins “putative targets of Pmk1”. Importantly, using this approach we identified some previously reported Pmk1 targets. For example, we observed Pmk1-dependent phosphosites at S126 and S158 in the transcription factor Hox7 (**Figure 5C and 5D**), consistent with a previous Hox7 study ^7^. Additionally, important components of the Pmk1 pathway, such as the MAPKKK Mst11, MAPKK Mst7 and adaptor protein Mst50 also showed Pmk1-dependent phosphorylation (**Figure 5C and 5D**). In total, our PRM analysis of a time series study of infection-related development in *M. oryzae* identified a set of 33 putative substrates of Pmk1.

**Figure 5.**
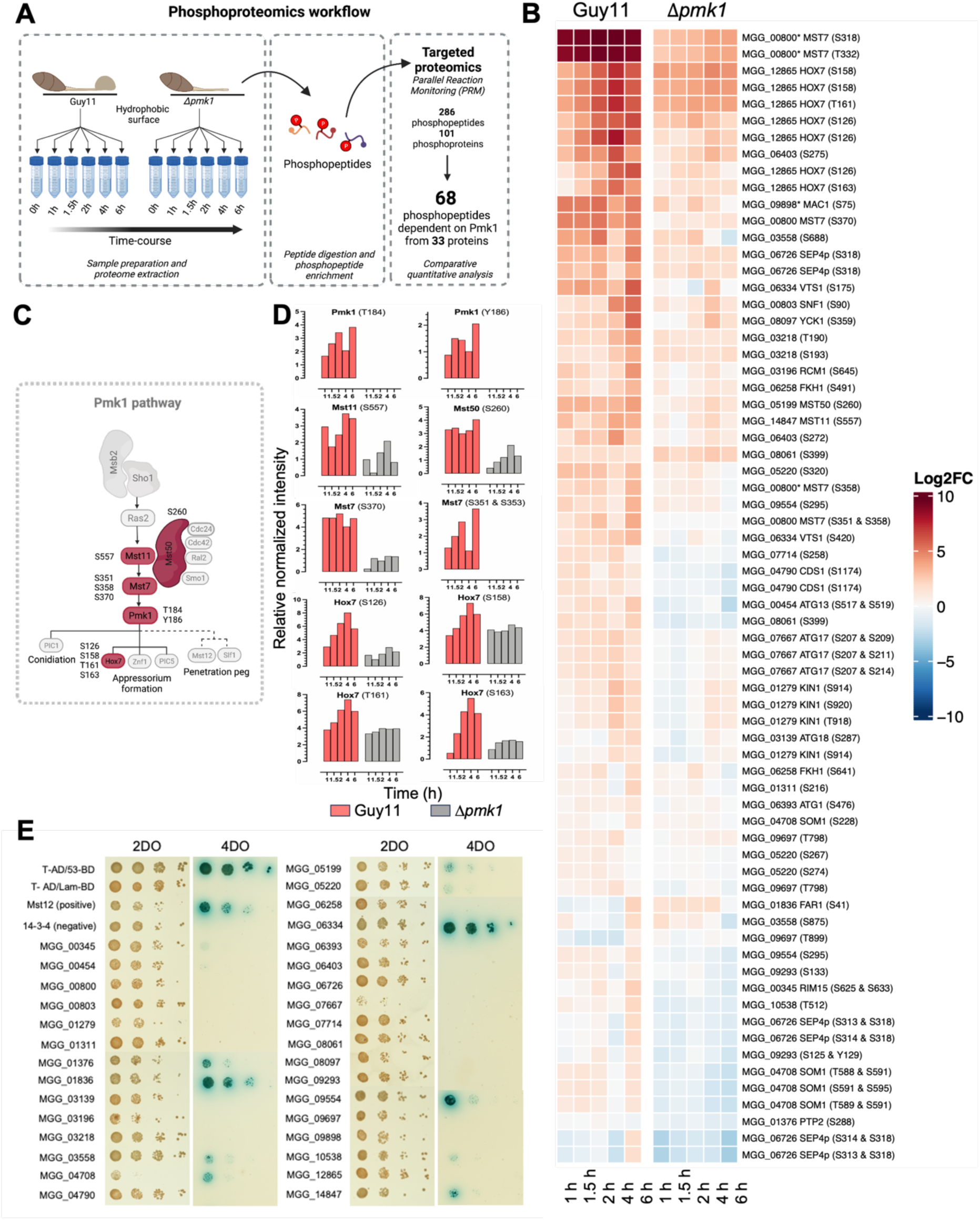
Quantitative phosphoproteomics defines 33 putative targets of the Pmk1 MAPK pathway. **(A)** Schematic to show the phosphoproteomic workflow for the quantitative analysis experiment between Guy11 and 11*pmk1* to determine Pmk1 direct targets in *M. oryzae*. Spores and germinated cells were synchronized at 0h, 1h, 1.5h, 2h, 4h and 6h. Phosphopeptides were enriched using TiO2 microspheres and were subjected to liquid chromatography-tandem mass spectrometry (LC-MS/MS). Samples were analyzed by Parallel Reaction Monitoring (PRM). **(B)** Heatmap to show the differentially phosphorylated phosphopeptides in Guy11 and 11*pmk1.* Each row corresponds to a single phosphosite. **(C)** Schematic representation of components of the Pmk1 signaling pathway in *M. oryzae*. Differentially phosphorylated residues in are indicated for each protein (adjusted p value <= 0.05). **(D)** Bar graphs to show relative normalized intensity determined by PRM of peptides associated to Mst11, Mst7, Pmk1, Mst50 and Hox7 during early infection from 0-6h in Guy11 and 11*pmk1*. **(E)** Yeast-two-hybrid (Y2H) assay to determine the interaction of Pmk1 with its putative direct targets. Protein interactions were tested in yeast grown on SD medium -Trp - Leu -Ade -His + alpha X-gal +Au (4DO panels). Viability of all transformed yeast cells was demonstrated by growth on SD medium - Trp -Leu (2DO panels). Yeast cells were inoculated onto media as a tenfold dilution series. Mst12 was used as the positive control.

The identified potential Pmk1 targets are broadly representative of cellular processes implicated in appressorium morphogenesis, based on previous studies ^7^, but have diverse functions. Based on the Magnagenes database of gene functional studies in *M. oryzae* ^34^, the function of 22 of the putative Pmk1 targets have already been studied in the blast fungus. However, 11 proteins of this subset have not yet been described (**Table 3**). Using gene ontology annotation, we assigned a function to each of these proteins where possible. From the proteins previously studied in *M. oryzae*, we found 5 kinases, 4 transcription factors, 1 transcriptional regulator and 1 phosphatase. Additionally, we also found 5 autophagy-related proteins, 5 components of the Pmk1 cascade, 2 components of the cAMP signaling pathway and 1 cytoskeleton related-protein. To determine which proteins might be direct targets of Pmk1, we carried out yeast-two-hybrid (Y2H) screening for these 33 putative targets using Pmk1 as bait on high stringency selection media (quadruple drop-out medium) (**Figure 5E**). We observed direct protein-protein interactions with Pmk1 in 9 of the 33 putative targets. Interactors include the transcription factors Far1 ^35^ implicated in the regulation of lipid metabolism associated with appressorium turgor generation and Som1, which links cell cycle control of appressorium morphogenesis with cAMP signaling ^36,37^, as well as Mst50 and Mst11, the phosphatase Ptp2, and a set of previously uncharacterized proteins, including a potential regulatory protein called Vts1. Taken together, these results show that quantitative comparative phosphoproteomics, can identify putative direct targets of the Pmk1 MAP kinase.

**Table 3.** Pmk1 putative targets identified during appressorium formation by PRM.

| Gene ID | Name | Function/ Process | Pmk1 dependent phosphorylation site |
| --- | --- | --- | --- |
| MGG_01311 | Nuclear elongation protein | Uncharacterized | S216 |
| MGG_03218 | Calcipressin protein | Uncharacterized | T190, S193 |
| MGG_03558 | PH domain protein | Uncharacterized | S666, S688, S875 |
| MGG_05220 | S/T protein kinase | Uncharacterized | S267, S274, S320 |
| MGG_06334 | Vts1 | Uncharacterized | S175, S420 |
| MGG_06403 | PH domain protein | Uncharacterized | S272, S275 |
| MGG_07714 | Actin cytoskeleton organization protein | Uncharacterized | S258 |
| MGG_09293 | Cell division control protein | Uncharacterized | S97, S125, Y129, S133 |
| MGG_09554 | Anaphase-promoting complex subunit | Uncharacterized | S295 |
| MGG_09697 | PH domain protein | Uncharacterized | T798, T899 |
| MGG_10538 | RSC complex subunit | Uncharacterized | T512 |
| MGG_00345 | RIM15 | Kinases | S402, S625, S633 |
| MGG_00803 | SNF1 | Kinases | S90 |
| MGG_01279 | KIN1 | Kinases | S914, T918, S920 |
| MGG_04790 | CDS1 | Kinases | S1174 |
| MGG_08097 | YCK1 | Kinases | S359 |
| MGG_06393 | Atg1 | Autophagy | S476, S547 |
| MGG_00454 | Atg13 | Autophagy | S517, S519, S920 |
| MGG_07667 | Atg17 | Autophagy | S207, S209, S211, S214 |
| MGG_03139 | Atg18 | Autophagy | S287, S295 |
| MGG_08061 | Atg28 | Autophagy | S399 |
| MGG_14847 | Mst11 | Pmk1 pathway | T551, S557 |
| MGG_00800 | Mst7 | Pmk1 pathway | S351, S358, S370 |
| XP_368444.2 | Mst7 | Pmk1 pathway | S318, T332, S358 |
| MGG_05199 | Mst50 | Pmk1 pathway | S260 |
| MGG_01836 | FAR1 | Transcription factor | S41 |
| MGG_06258 | FKH1 | Transcription factor | S491, S641 |
| MGG_12865 | HOX7 | Transcription factor/<br>Pmk1 pathway | S126, S158, T161, S163 |
| MGG_04708 | SOM1 | Transcription factor/<br>cAMP pathway | S228, T588, T589, S591, S595 |
| MGG_09898 | MAC1 (XP_365053.1) | cAMP pathway | S75 |
| MGG_06726 | Septin4 | Cytoskeleton related | S313, S314, S318 |
| MGG_03196 | RCM1 | Transcriptional regulator | S645 |
| MGG_01376 | PTP2 | Phosphatase | S288 |

### Vts1 is a novel component of the Pmk1 MAPK pathway during rice blast disease

To test the validity of our quantitative phosphoproteomic approach, we decided to investigate whether Vts1 is a direct phosphorylated target of Pmk1 during appressorium development. To do this we first tested whether Pmk1 associated with Vts1 *in vivo*. We found that Vts1 is able to interact strongly with Pmk1 in a stringent Y2H assay (**Figure 5E and 6A**). To show whether this interaction occurred during appressorium development, we carried a co-immunoprecipitation experiment in appressorial protein extracts. We found that Pmk1 associates with Vts1 after 4h, during appressorium development, suggesting a possible role during development (**Figure 6B**). From PRM analysis, we identified 3 phosphorylation sites in Vts1 (**Figure 6D**). However, only S175 and S420 are differentially phosphorylated in the presence of Pmk1 in our PRM analysis (**Figure 6E**). To differentiate between direct and indirect effects of the absence of Pmk1 activity, we used a Pmk1 analogue-sensitive mutant (*pmk1^AS^*) ^6^, which can be inhibited selectively by addition of the ATP analogue 1-Napthyl-PP1. In this way, we were able to inhibit Pmk1 *in vivo* during early appressorium formation (**Figure 6F and S6**). We used PRM to accurately measure phosphopeptide abundance, which showed that specific inhibition of Pmk1 at 1-4h post-germination, affects phosphorylation at S175 and S420 of Vts1 but not at T14 (**Figure 6F**). To test whether Pmk1 phosphorylates Vts1 at S175 and S420 we carried out an *in vitro* kinase assay, using recombinant Vts1 and Pmk1. We used a recombinant constitutively active MAPKK from tobacco (Nicotiana tabacum) NtMEK2^DD^ to activate Pmk1 as previously shown ^7,91^ (**Figure S7**). The *in vitro* kinase assay showed that Pmk1 specifically phosphorylates Vts1 in a [S/T]P motif (**Figure 6C and S7D)** and mass spectrometry demonstrated that S175 and S420 are indeed phosphorylated by Pmk1 (**Figure S7E**). These results indicate that Pmk1 can associate with and specifically phosphorylate Vts1 at residues S175 and S420, consistent with Vts1 being a direct substrate of Pmk1.

**Figure 6.**
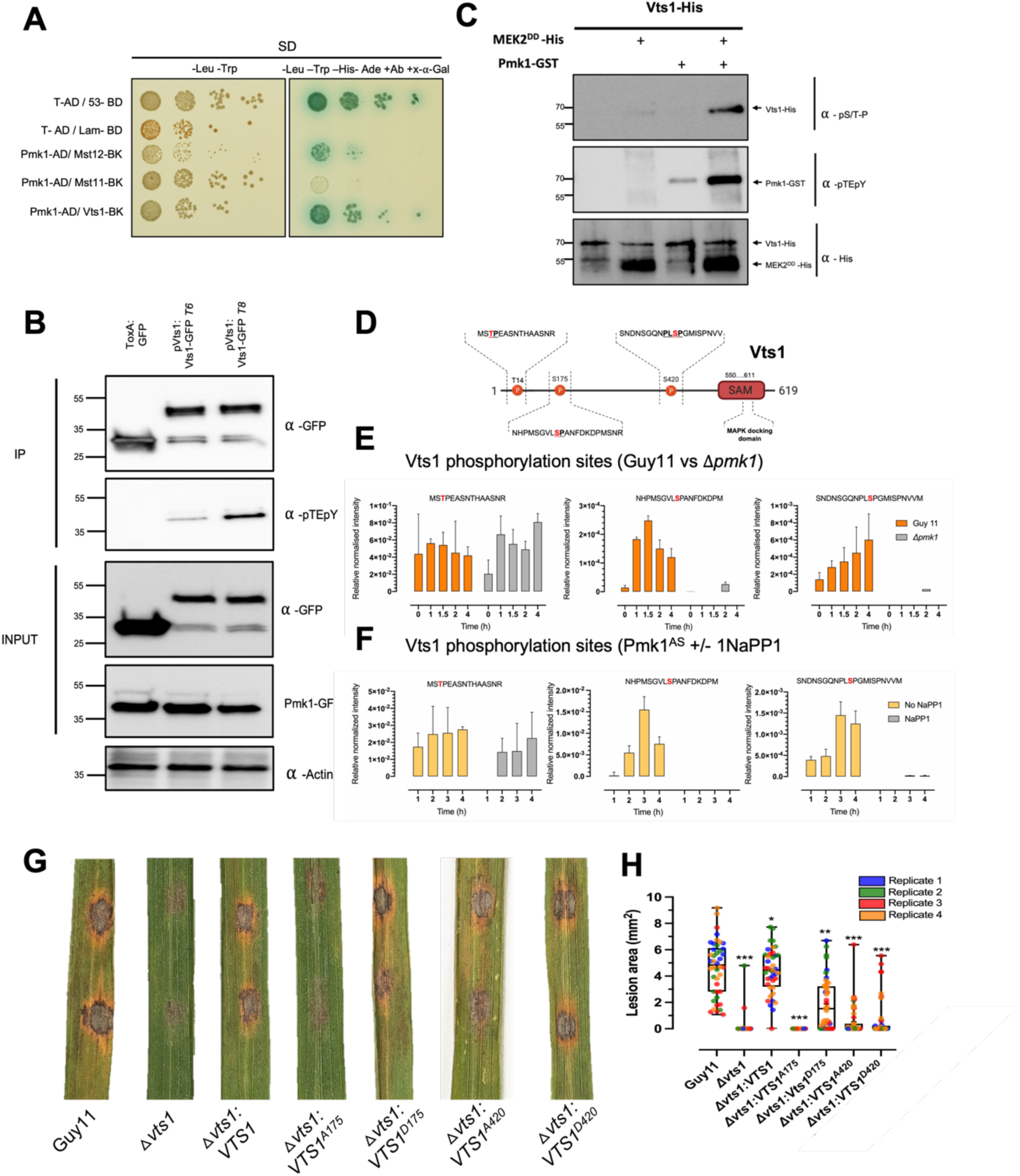
Vts1 is a novel target of the Pmk1 MAPK pathway necessary for rice blast disease. **(A)** Yeast-two-hybrid (Y2H) assay to determine the interaction of Pmk1 with its putative direct targets. Protein interactions were tested in yeast grown on SD medium -Trp - Leu -Ade -His +X gal +Au (4DO panels). Viability of all transformed yeast cells was demonstrated by growth on SD medium -Trp -Leu (2DO panels). Yeast cells were inoculated onto media as a tenfold dilution series. Mst12 was used as the positive control. **(B)** Co-immunoprecipitation of Vts1-GFP. C-terminal GFP tagged Vts1 was transformed into *M. oryzae* Guy11. Anti-pTEpY antiserum was used to detect double phosphorylated Pmk1. Immunoprecipitates obtained with anti-GFP antiserum, and total proteins extracts, were probed with appropriate antisera. **(C)** Western blot analysis of *in vitro* phosphorylation experiment between Pmk1 and Vts1 (N- terminally tagged with 6xHis). Proteins were immunoblotted with appropriate antisera (listed on the right). Arrows indicate expected band sizes. **(D)** Schematic diagram to show Vts1 phosphorylated residues T14, S175, S420 and position of the predicted SAM domain. **(E-F)** Relative normalized intensity determined by PRM of Vts1 phosphopeptides associated with T14, S175 and S420 during appressorium development from 0-4h in *M. oryzae* for **(E)** Guy11 and the 11*pmk1* mutant, and **(F)** the analogue-sensitive mutant *pmk1^AS^* in the presence or absence of the inhibitor 1NaPP1. **(G)** Leaf drop assay using 3-week-old seedlings of rice cultivar CO-39 that were inoculated with equal amounts of conidial suspensions of Guy11, 11*vts1* and Vts1 phosphorylation mutant strains (10^5^ conidia mL^-1^) in 0.2% gelatin. Seedlings were incubated for 5 days to develop blast disease at 26 °C. Fully susceptible, sporulating disease lesions can be distinguished by their grey centres. **(H)** Bar chart to show the lesion size for each *M. oryzae* phosphoallele. \**P* < 0.05: \*\**P* < 0.01; \*\*\**P* < 0.001 represent significance using unpaired two-tailed Student’s *t*-test. Data from four biological replicates.

To investigate the role of Vts1 in fungal pathogenicity, we generated a targeted deletion mutant of *VTS1* in *M. oryzae* Guy11 (**Figure S8 and S9A**). We observed that 58% of Δ*vts1* appressoria show aberrant development compared to Guy11 (**Figure S9B**) and Δ*vts1* mutants were severely impaired in their ability to cause rice blast disease (**Figure S9C and S9D**). To study the role of Pmk1-dependent phosphorylation of Vts1, we generated phosphomimetic and non-phosphorylatable (‘phosphodead’) alleles of *VTS1* and transformed these into the Δ*vts1* mutant. We observed that appressorium formation was impaired only by the strain complemented with the phosphodead allele *VTS1*^A175^ but not by any of the other phosphorylation mutant variants (**Figure S9E**). Furthermore, leaf sheath and leaf drop infection assays showed that *VTS1*^A175^ does not produce rice blast disease lesions and infection progression is severely impaired (**Figure 6G and 6I)**. This provides strong evidence that Pmk1-dependent phosphorylation of S175 of Vts1 is necessary for development rice blast disease. Consistent with the requirement of S175 for fungal virulence, we found that Vts1 S175 is conserved among filamentous fungi, while S420 is not conserved in other fungal species (**Figure S10**). This suggests that S175 and S420 fulfil different distinct functions. When considered together, these results demonstrate that quantitative phosphoproteomics has the capacity to identify completely novel regulators of fungal virulence and enable the functional characterization of signaling pathways that govern plant infection by pathogenic fungi.

## DISCUSSION

Fungal pathogenicity is a complex phenotype that encompasses the ability of many fungal pathogens to develop specialized infection structures to breach the tough outer layers of plants, insects or human cells, to colonize living host tissue, to suppress immunity by deployment of large families of effector proteins, and finally to kill host cells and produce new infective propagules to infect new hosts. Very few global regulators of fungal pathogenesis have been identified to date, but MAP kinases appear to be widely conserved across fungi and in pathogenic species, they have been shown to control invasive growth and virulence (**Table 1**). The rice blast fungus is one of the most devastating pathogens in the world and the MAP kinase Pmk1 is a master regulator of infection-related development, regulating appressorium morphogenesis ^5^, appressorium function ^7,33,38^ and invasive growth in plant tissue ^6^. This single kinase has been shown to control 49% of total gene expression during appressorium development, suggesting a very broad role in the control of fungal development and physiology^7^. Identifying the exact targets of master-regulator kinases across pathogenic fungi and understanding the degree of conservation between them could provide new information that can be exploited to control of some of the most devastating diseases across the world.

In this study we aimed to test whether quantitative phosphoproteomics could provide new insight into the biology of infection by pathogenic fungi and identify downstream processes via its direct phosphorylation substrates. The first major conclusion of this study is that the phosphorylation landscape of fungal infection by the rice blast fungus is complex and highly dynamic. Dramatic changes in phosphorylation occur as early as initial contact to an inductive surface and extend to appressorium differentiation. We found a total of 8005 peptides corresponding to 2062 proteins change in abundance and/or are differentially phosphorylated in just 6h of development. The approach revealed that critical physiological processes previously reported to be essential for infection are highly dynamic and tightly regulated by phosphorylation. These processes include autophagy, which is essential for recycling the contents of the three-celled conidium into the unicellular appressorium ^20^, lipid metabolism ^38,39^, which is essential for glycerol synthesis that acts as the compatible solute in an appressorium for generation of its enormous turgor, and melanization which is essential for the appressorium to generate pressure ^40,41^. Many other processes including cell cycle regulation, cell wall biogenesis, intracellular trafficking, secondary metabolism are also clearly regulated by dynamic changes in phosphorylation during infection.

The second major conclusion of the study is that patterns of phosphorylation have been conserved across diverse fungal species, revealing phosphorylation signatures correlated with fungal pathogenesis and infection structure development, many of which are likely to be dependent on Pmk1-related MAPK activity. Comparative analysis with 41 fungal species provided evidence that elements of the phosphorylation landscape identified in *M. oryzae* are conserved across different fungal species. A sub-set of phosphorylation sites are, for example, highly conserved in plant-associated fungal species, with further phosphosites conserved only in fungal pathogens that invade cereal hosts, and others only present in fungal pathogens that elaborate melanised force-generating appressoria, like the blast fungus. These phosphosites have the potential to enable mining of conserved physiological processes associated with fungal pathogenesis, including infection structure development, and the invasion of host tissue. Specific processes regulated by patterns of phosphorylation, for example, in *Colletotrichum* and *Magnaporthe* include trehalose and glycogen metabolism, regulated lipolysis, cytoskeletal re-modelling and BAR domain proteins implicated in membrane curvature generation. These make sense in the context of development of a melanized, high pressurized appressorium, but have not been studied in a comparative way between these species before.

Quantitative phosphoproteomic analysis enabled the detailed analysis of Pmk1-dependent phosphorylation in *M. oryzae*, revealing the specific signaling pathways targeted by the MAPK. These include the cAMP-dependent protein kinase A pathway, which serves a role both in surface sensing and the physiological regulation of compatible solute generation in the appressorium ^1,2^ and the regulators of autophagy, such as the Atg1 kinase, Atg13 and Atg17, which initiate phagophore formation at the onset of autophagy ^42^. This link is consistent with previous studies in which autophagy was shown to be impaired in a *Δpmk1* mutant ^33^, but also show the likely mechanism of by which Pmk1 exerts this control. Completely new insights have also been provided, however, such as the link with the Sln1 turgor sensor kinase, which is necessary to regulate polarized growth and penetration peg emergence once a threshold of turgor has been generated ^3,4^. Pmk1 is necessary for phosphorylation of the Sln1 histidine kinase and an interacting stretch-activated gated ion channel protein Mic1, as well as the components of the cell integrity pathway, such as protein kinase C that are necessary for regulating the changes in cell wall structure associated with the resumption of polarized growth. Interestingly, the NADPH oxidase Nox2, which is necessary for septin aggregation at the appressorium pore is also phosphorylated in a Pmk1-dependent manner, along with its Bem1 regulator. Similarly, the newly identified Vast1 pathway ^43^, which is necessary for TOR-dependent plasma membrane homoeostasis is also regulated in a Pmk1-dependent manner. This is also consistent with aminophospholipid regulators, Pde1 ^44^ and Apt2 ^28^, which are both necessary for appressorium function, being phosphorylated in a Pmk1-dependent manner. A third major conclusion of our study is therefore that quantitative phosphoproteomics can provide unparalleled insight into the regulatory processes controlled by Pmk1 during infection.

To test the ability to identify and characterize direct substrates of the Pmk1 MAPK, we focused on investigating a phosphorylated Pmk1 interactor called Vts1, a protein of unknown function in the rice blast fungus. We observed that Vts1 contains a sterile alpha motif (SAM) domain. SAM domain-containing proteins have been previously reported as important regulators of MAPK signaling cascades (Kim & Bowie, 2003) and proteins containing SAM domains are versatile because this domain has documented to take part in various interactions. They can, for example, show binding affinity to other SAM and non-SAM domain proteins, but can also show binding affinity to lipids and RNA ^46^. Importantly, SAM domains have been reported to mediate associations of MAPK modules in different fungi. In *Schizosaccharomyces pombe*, for instance, association between Ste4 and Byr2 occurs via a SAM motif ^47^. Similarly, in *S. cerevisiae*, the interaction between Ste11 and Ste50 is mediated by a SAM domain ^48^. In *M. oryzae*, the MAPKK Mst11 and the putative scaffold protein Mst50 in the Pmk1 pathway also both contain SAM domains ^10^. Furthermore, it has been demonstrated that Mst50-Mst11 interaction occurs via their respective SAM domains and that this is essential for appressorium development and plant infection ^11^. In this study, we have provided evidence that Vts1 is phosphorylated directly by Pmk1 based on parallel reaction monitoring, the analysis of a conditional analogue-sensitive Pmk1 mutant, and an *in vitro* kinase assay. Furthermore, Vts1 is important for fungal virulence and necessary for correct appressorium morphogenesis. Finally, we demonstrated that a single Vts1 phosphosite S175 is necessary for its biological activity during plant infection. These results therefore validate the use of phosphoproteomics as a means of identifying new determinants of pathogenicity in the blast fungus and thereby revealing how Pmk1 exerts such a major role in the regulation of fungal infection.

When considered together, this study demonstrates the importance of phosphoproteomic changes during infection-related development by a pathogenic fungus. The conservation of nearly 1200 phosphorylated residues in a group of 41 fungal species also reveals proteins required for core functions as well as potential phosphoproteins associated with a plant pathogenic lifestyle. This underscores the potential of our phosphoproteome data sets as a resource that can be mined by the fungal research community to identify novel virulence determinants in a wide range of fungal species.

## ACKNOWLEDGMENTS

We thank Prof. Sophien Kamoun for critical input on this manuscript. We thank the John Innes Centre (JIC) Bioimaging staff for providing technical support on SEM images. We thank Dr. Iris Eiserman for providing the Septin4 pGBKT7 construct. This work was supported by grants to NJT from the Gatsby Charitable Foundation and Biotechnology and Biological Sciences Research Council (BBSRC) BBS/E/J/000PR9797 and a BBSRC grant to NJT and FLHM (BB/V016342/1). NCM was supported by a John Innes Foundation, Sainsbury Laboratory

Rotation Programme PhD Studentship and Consejo Nacional de Humanidades, Ciencia y Tecnologías (CONAHCYT).

## AUTHORS CONTRIBUTIONS

NJT and FLHM conceived the project. NJT and FLHM guided the execution of the experiments and oversaw the project. NCM, MOR, PD, CJ, LR, MJB, AEB, JS, BT, XY, WM, VW, DM and FLHM did the experiments and analyzed the data. NCM prepared figures and tables. NCM, NJT, and FLHM wrote the manuscript with contributions from all authors.

## KEY RESOURCES TABLE

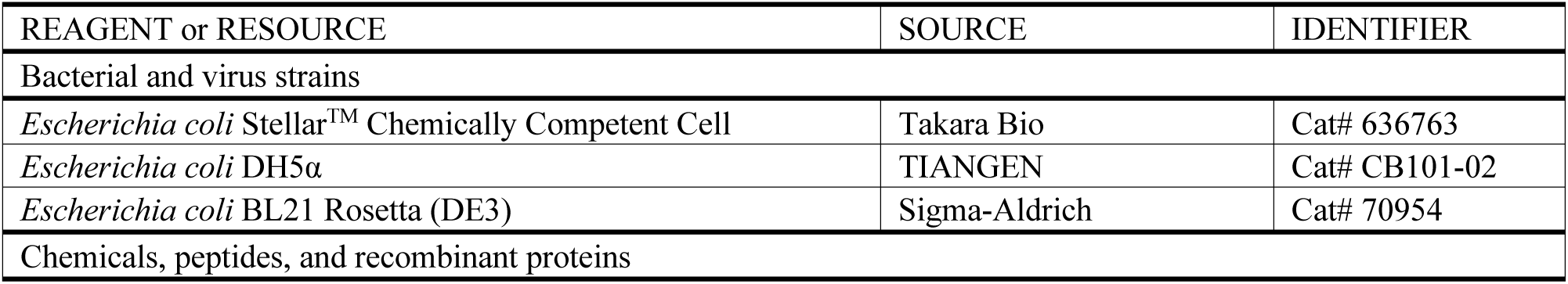
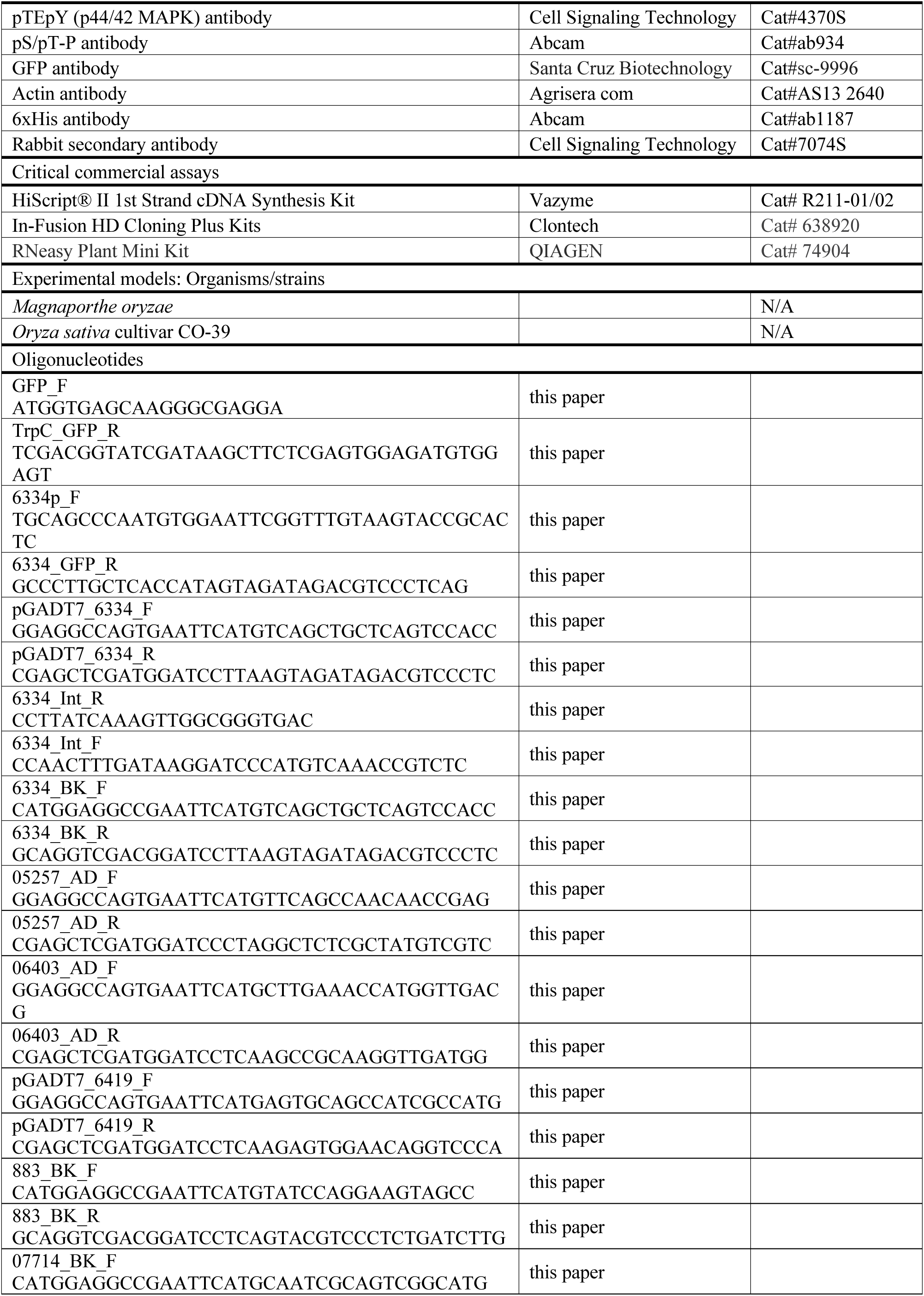
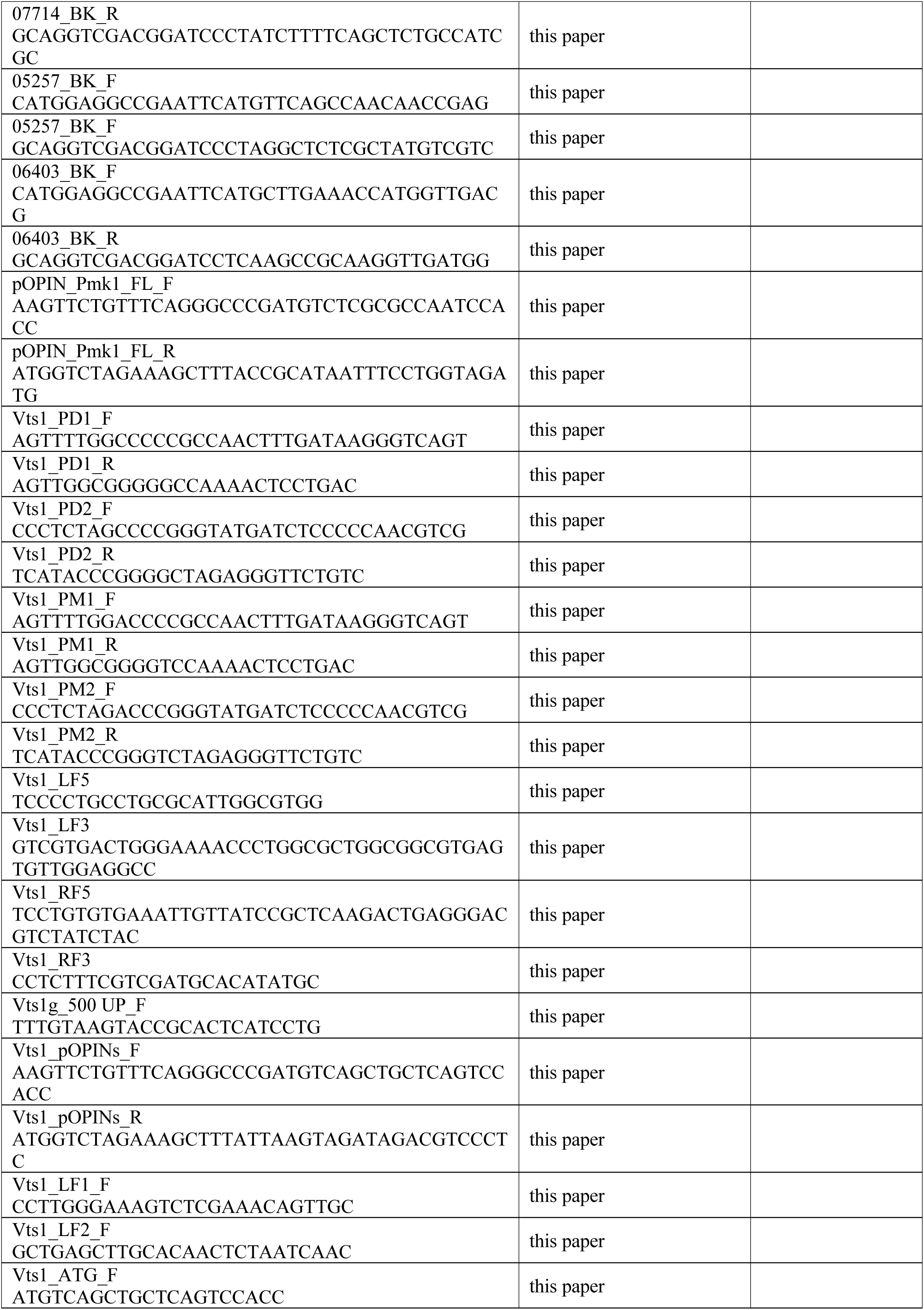
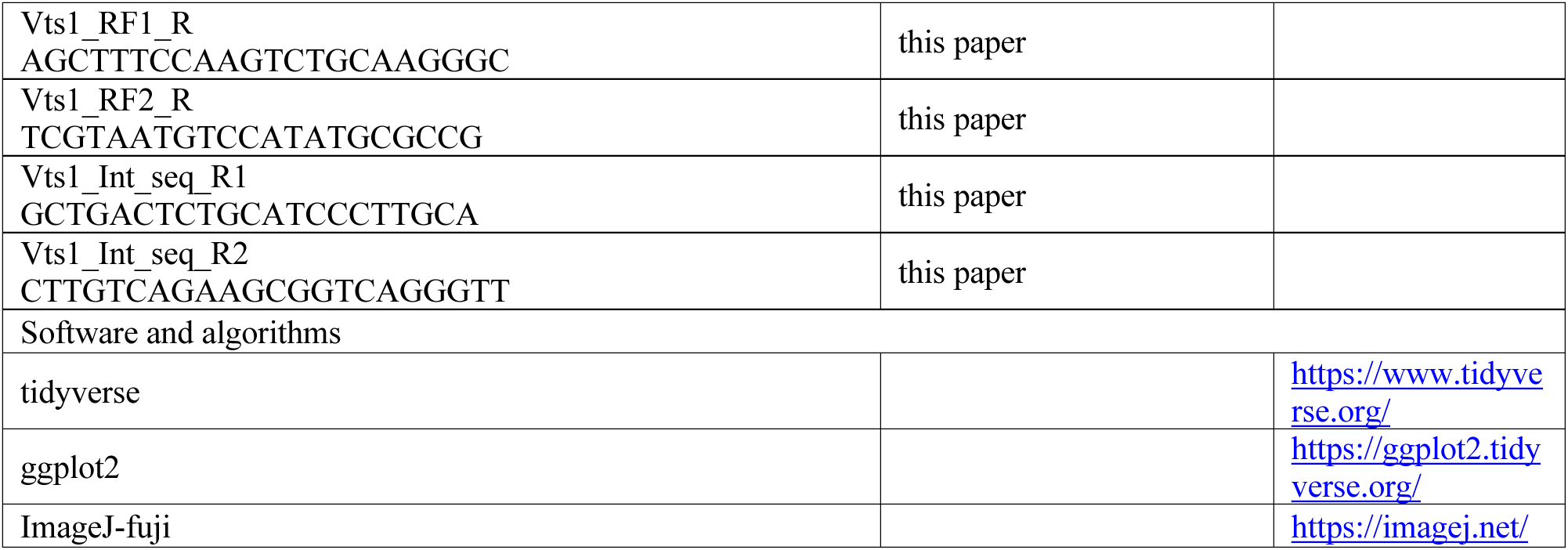

## RESOURCE AVAILABILITY

### Lead Contact

Frank Menke is the lead contact for MS-related data and proteomics and Nicolas Talbot is the lead contact for biological materials and fungal strains.

### Materials Availability

The DDA proteomics data have been deposited to the ProteomeXchange Consortium via the PRIDE ^82^ partner repository with the dataset identifier PXDxxxxx and xxxxxxxxxxx. and targeted proteomic data via Panorama ^83^.

## EXPERIMENTAL MODEL AND SUBJECT DETAILS

### Fungal strains

*Magnaporthe oryzae* strains used in this study were routinely grown on agar plates with solid complete medium. Fungal strains were incubated at 26°C with a 12 h light and dark cycle ^84^. For long-term storage, *M. oryzae* strains were grown over sterile filter paper discs (Whatman International) placed on complete medium agar plates. The paper discs were then dehydrated and stored at -20°C.

### Rice plants

Blast susceptible rice (*Oryza sativa*) cultivar CO-39 plants were used in this study. Plants were grown in controlled environment rooms at 26°C day temperature and 24 °C night temperature, 16-hour light period, and 85% humidity ^84,85^.

## METHODS DETAILS

### *M. oryzae* appressorium in vitro assay for proteomics analysis

Conidia were harvested from a Petri dish culture using a sterile disposable plastic spreader in 3 mL sterile distilled water from 8–12 days old cultures grown on CM agar. The conidial suspension was filtered through sterile Miracloth (Calbiochem, UK) and fractionated by centrifugation at 5000 x *g* (Beckman, JA-17) for 15 min at room temperature. The pellet of conidia was re-suspended in 0.2 % (w/v) gelatin (BDH) and the spore concentration determined using a haemocytometer (Improved Neubauer, UK). Spores were diluted to a final concentration of 5 x10^4^ conidia mL^-1^. Conidia were quantified and then diluted in sterile water to 7.5×10^5^ conidia/mL in the presence of 50 ng/µL 1,16-Hexadecanediol (Sigma SA). For microscopic observations, a 50 μL aliquot of conidial suspension was inoculated onto a borosilicate glass coverslip (Menzel-Gläser, Fisher Scientific UK Ltd.) and placed on a moist paper towel. Conidia were incubated at 24 °C and observed as indicated. For large-scale conidial germination assays, conidial suspensions were poured into square petri plates (12 cm X 12 cm X 1.7 cm) (Greiner Bio One) to which 10 glass cover slips (Menzel-Gläser, Fisher Scientific UK Ltd.) were attached by adhesive. Appressorium formation was monitored under a Will-Wetzlar light inverted microscope (Wilovert^®^, Hund Wetzlar, Germany) for ensuring homogeneous and synchronized infection structure formation. Samples were collected as indicated by scraping the surface of the coverslips with a sterile razor blade (Fisher Scientific, UK). Harvested samples were immediately frozen in liquid nitrogen and stored at −80 °C for subsequent protein extraction. The appressorium *in vitro* development assay was adapted from Hamer et al., 1988 ^86^.

### Virulence analysis of fungal strains on rice

Conidia were harvested from a Petri dish culture using a sterile disposable plastic spreader in 3 mL sterile distilled water from 8–12 days old cultures grown on CM agar. The conidial suspension was filtered through sterile Miracloth (Calbiochem, UK) and fractionated by centrifugation at 5000 x *g* (Beckman, JA-17) for 15 min at room temperature. The pellet of conidia was re-suspended in 0.2 % (w/v) gelatin (BDH) and the spore concentration determined using a hemocytometer (Improved Neubauer, UK). Spores were diluted to a final concentration of 5 x10^4^ conidia mL^-1^. For spray infection assays, the spore suspension was used to infect rice using an airbrush (Badger. USA). After spray inoculation, the plants were covered in polythene bags and incubated in a controlled plant growth chamber (Conviron, UK) at 24°C for 48 h with a 12 h light and dark cycle, and 85% relative humidity. The inoculated plants were incubated for 5-6 days before scoring the lesions ^87^. For leaf drop assays, the spore suspension was drop-inoculated on detached rice leaves using a micropipette. Rice CO-39 plants were grown for 3 weeks in 9 cm diameter plastic plant pots.

### Confocal laser scanning microscopy

Confocal laser scanning fluorescence microscopy was performed on a Leica TCS SP8 microscope using 40x or 63x/1.4 oil immersion objective lens. Images were acquired using Leica LAS AF software (Leica Macrosystems Inc., Buffalo Grove, IL, USA). Fluorescence was observed using HyD detectors and white laser. The filter sets used for GFP were excitation wavelength 488 nm and emission collected at 495–550 nm. For RFP probes, the excitation wavelength was 543 nm and emission collected at 584 nm. Confocal microscopy images were processed with the Leica LAS AF software and ImageJ (2.0) programs.

### Cryo-Scanning Electron Microscopy

The infected rice leaf samples were mounted on an aluminium stub using Tissue Tek^R^ (BDH Laboratory Supplies, Poole, England). The stub was then immediately plunged into liquid nitrogen slush at approximately -210°C to cryo-preserve the material. The sample was transferred onto the cryo-stage of an ALTO 2500 cryo-transfer system (Gatan, Oxford, England) attached to an FEI Nova NanoSEM 450 (FEI, Eindhoven, The Netherlands). Sublimation of surface frost was performed at -95°C for three minutes before sputter coating the sample with platinum for 3 mins at 10mA, at colder than -110°C. After sputter-coating, the sample was moved onto the cryo-stage in the main chamber of the microscope, held at approximately -125°C. The sample was imaged at 3kV and digital TIFF files were stored.

### Yeast-Two-Hybrid (Y2H) analysis

Desired constructs in pGBKT7 and pGADT7 were co-transformed into chemically competent *Saccharomyces cerevisiae* Y2HGold cells (Takara Bio, USA) using the commercial kit Frozen-EZ Yeast Transformation II^TM^ (Zymo Research, UK) as detailed in the user manual. The Matchmaker^®^ Gold Yeast Two-Hybrid System (Takara Bio USA) was used to detect protein– protein interactions between Pmk1 and its putative targets. Single co-transformed colonies grown on selection plates were inoculated in 5mL of SD^-Leu-Trp^ and grown overnight at 30°C. Saturated culture was then used to make serial dilutions of OD_600_ 1, 1^-1^, 1^-2^, 1^-3^, respectively. An aliquot of 5μl of each dilution was then spotted on a SD^-Leu-Trp^ plate as a growth control, SD^-Leu-Trp-His^ (low stringency media) and SD^-Leu-Trp-Ade-His^ (high stringency media) plate containing X-α-gal and aureobasidin A. Plates were imaged after incubation for 60 - 72 hr at 30°C.

### *M. oryzae* genomic DNA purification

For large-scale DNA extraction, fungal mycelium was generated by growing fungal culture on either cellophane discs or liquid as previously described ^85^. Using a mortar and pestle, 7-12 days old mycelium was ground into powder. Mycelial powder was decanted to a 1.5 mL microcentrifuge tube and mixed with 500 μL of pre-warmed CTAB (2% (w/v) Hexadecyltrimethylammonium Bromide (CTAB), 100 mM Tris base, 10 mM EDTA and 0.7 M NaCl) and incubated at 65°C with gentle mixing every 10 min. An equal volume of chloroform iso-amyl alcohol (CIA) was added, mixed thoroughly, and incubated with shaking for 30 min at room temperature. This was followed by centrifugation at 17000 x *g* for 10 min. This step was repeated twice by adding equal volumes of CIA and mixing vigorously on a shaker before centrifugation. The final supernatant was transferred into a clean sterile microcentrifuge tube and of isopropanol (2 x vol) added before incubating at -20°C overnight. The samples were centrifuged at 17000 x *g* for 10 min and the supernatant (isopropanol) was gently removed and the resulting pellet re-suspended in 500 μL sterile distilled water (SDW) and left to dissolve at room temperature with gentle tapping to mix. Sodium acetate (NaOAc) (0.1 vol) and 100% ethanol (2 vol) were added to re-precipitate nucleic acids. The mixture was incubated at -20°C for 2 h and pelleted by centrifugation at maximum speed, before washing with 400 μL of 70% (v/v) ethanol. The DNA was re-suspended in nuclease-free water. RNase (2 μL) was added and incubated at 37°C for 1 h to digest contaminating RNA.

### Southern blotting

In this study, Southern blot analysis was used to determine positive *M. oryzae* null mutants for the *VTS1* gene. DNA digestion of *M. oryzae* transformants was performed overnight using *Hind*III endonuclease and subsequently fractionated by electrophoresis in an agarose gel at 100V. Fragments of genomic DNA were separated in agarose gels were transferred to Hybond-NX (Amersham Biosciences). Prior to blotting, partial depurination of DNA molecules was performed to enhance DNA transfer by submerging the agarose gel in 0.25 M with gentle rocking. Gels were then neutralized by replacing HCl with 0.4 M NaOH. For transfer of DNA from the agarose gel to the positively charged membrane, blots were carried out using a 0.4M NaOH transfer buffer that was drawn through a wet paper wick (Whatman /international) supported by a Perspex panel onto which the agarose gel was placed. A sheet of Hybond-NX membrane was then laid on the gel and positions of the wells were pencil marked. Three layers of Whatman 3MM paper and a stack of paper towels (Kimberley Clark Corporation) were laid over the membrane followed by a glass plate and a 500 g weight were placed on the stack as a weight. The transfer was left at room temperature overnight. Then, the nucleic acid was fixed to the membrane by UV crosslinking to the membrane with 120 milijoules.cm^-2^ using a BLX crosslinker (Bio-Link).

### *M. oryzae* whole genome sequencing

Purified DNA was obtained using the CTAB procedure, as previously described ^85^. A NanoDrop spectrophotometer (Thermo Scientific, UK) and a Qubit BR assay (Thermo Fischer, USA) were used to analyze template quality and determine the concentration of double-stranded DNA. Sequencing was carried out using Novogene Sequencing services (Cambridge, UK). Whole genome sequencing was performed on NovaSeq 6000 system (Illumina), with two lanes per sample. BAM files were created by using bowtie2 for aligning raw reads to the *M. oryzae* reference genome 70-15. Finally, the IGV viewer was used for the visualization of the generated BAM files.

### Protoplast-mediated transformation of *M. oryzae*

A section of 2.5 cm^2^ mycelium from a *M. oryzae* plate culture (8-10 days-old) was blended in 150 mL CM liquid and incubated at 25°C, shaking (125 rpm) in an orbital incubator for 48h. Fresh ST (sucrose, 0.6M, Tris-HCl 0.1 M (pH 7), STC (sucrose, 1.2 M, Tris-HCl, 10 mM (pH 7.5)) and PTC (PRG 4000, 60%, Tris-HCl, 10 mM (pH 7.5), calcium chloride) buffers were prepared and stored at 4°C. The culture was harvested by filtration through sterile Miracloth and the mycelium washed with sterile deionized water (SDW). The mycelium was transferred to a 50 mL falcon tube with 40 mL OM buffer (1.2 M magnesium sulfate, 10 mM sodium phosphate (pH5.8), Glucanex 5% (Novo Industries, Copenhagen)). Mycelium in the falcon tube with OM buffer was shaken gently to disperse hyphal clumps. Then, it was incubated at 30°C with gentle (75 rpm) shaking, for 3 h. The digested mycelium was transferred to two sterile polycarbonate Oakridge tubes (Nalgene) and overlaid with an equal volume of cold ST buffer. Resulting protoplasts were recovered at the OM/ST interface by centrifugation at 5000 x *g*, for 15 min at 4°C in a swinging bucket rotor (Beckman JS-13.1) in a Beckman J2.MC centrifuge. Protoplasts were recovered and transferred to a sterile Oakridge tube, which was then filled with cold STC buffer. The protoplasts were pelleted at 3,000 x *g* for 10 min (Beckman JS-13.1 rotor). This wash was carried out twice more with STC, with complete re-suspension of the pellet After the last wash, protoplasts were resuspended in 1 mL of STC and checked by microscopy. In an Eppendorf tube, an aliquot of protoplasts was combined with 6 µg DNA. The mixture was incubated at room temperature for 30 min. After incubation, 1 mL of PTC was added in 2 aliquots, mixed gently by inversion, and incubated at room temperature for 20 min. The transformation mixture was added to 150 mL of molten agar medium and poured into 5 sterile Petri dishes. For selection of transformants on hygromycin B (Calbiochem), plate cultures were incubated in the dark for at least 16 h at 24°C and then overlaid with approximately 15 mL of OCM/1% agar (CM osmotically stabilised with sucrose, 0.8M) containing 600µg mL^-1^ hygromycin B. For selection of bialophos (Basta) resistant transformants, OCM was replaced with BDCM (yeast nitrogen base without amino acids and ammonium sulfate, 1.7 g L^-1^ (Difco), ammonium nitrate, 2 g L^-1^ asparagine, 1 g L^-1^ glucose, 10 g L^-1^ sucrose, 0.8 (pH 6)). In the overlay, CM was replaced by BDCM without sucrose and hygromycin B was replaced by glufosinate (30 µg mL^-1^) from a stock at 100 mg mL^-1^ in DSW. For selection of sulfonylurea resistant transformants, OCM was replaced with BDCM and in the overlay, hygromycin B was replaced with chlorimuron ethyl, at 30 µg mL-1 freshly diluted from a stock solution, at 100 mg mL^-1^.

### SDS-PAGE and Western blot

Western blot analysis was performed on recombinant proteins and *M. oryzae* total protein. *M. oryzae* total protein samples were collected at indicated time point and snap-frozen in liquid-nitrogen. Lyophilised samples were lysed, and proteins were extracted with GTEN buffer (10 % glycerol, 25 mM Tris pH 7.5, 1 mM EDTA, 150 mM NaCl) with 10 mM DTT, 1% NP-40 and protease inhibitor cocktail (cOmplete™, EDTA-free; Merck), phosphatase inhibitor cocktail 2 (SigmaAldrich; P5726) and phosphatase inhibitor cocktail 3 (Sigma-Aldrich; P0044). After centrifugation at 13,000 rpm for 10 mins, protein concentration was measured and normalised with the Bradford assay (Protein Assay Dye Reagent Concentrate; Bio-Rad). After normalization, extracts were heated in 2× TruPAGE™ LDS Sample Buffer (SigmaAldrich) at 70 °C for at least 5 mins. Different percentage SDS-PAGE gels were used to run samples of difference sizes. Proteins were separated by SDS-PAGE and transferred onto a polyvinylidene diflouride (PVDF) membrane using a Trans-Blot turbo transfer system (Bio-Rad, Germany). The membrane was blocked with 3% BSA in Tris-buffered saline and Tween 20. Membranes were immunoblotted with antibodies specified in Table 2.1. Membrane imaging was carried out with an ImageQuant LAS 4000 luminescent imager (GE Healthcare Life Sciences, Piscataway, NJ, U.S.A.).

### Recombinant proteins production and purification

Recombinant pOPIN plasmids encoding 6xHisGST-Pmk1 and 6xHis-Vts1 were transformed into *E. coli* RosettaTM (DE3) cells. The bacteria were pre- inoculated in 100 mL of LB medium with carbenicillin and chloramphenicol overnight. An amount of 25 mL culture was then diluted into 1 L of autoinduction media (AIM) (10 g/L tryptone, 5 g/L yeast extract, 3.3 g/L (NH_4_)_2_SO_4_, 6.8 g/L KH2PO4, 7.1 g/L Na2HPO4, 0.5 g/L glucose, 2 g/L α- lactose, 0.15 g/L MgSO_4_ magnesium sulphate and 0.03 g/L trace elements) (Studier, 2005) with appropriate antibiotics and grown in at 37 °C (30 °C for Shuffle cells) for 6 h and then 16 °C overnight. Cells were harvested and resuspended in ice-cold lysis buffer (50 mM Tris-HCl pH 8.0, 50 mM glycine, 5% glycerol, 500 mM NaCl and 20 mM imidazole, supplemented with cOmpleteTM EDTA-free Protease Inhibitor Cocktail). The cells were then disrupted by sonication using a Vibra-CellTM sonicator (SONICS) with a single 13 mm probe, with the cells chilled on ice. The sonicator was set at 40 % amplitude, with a 1 s pulse followed by a 3 s pause, for 16 min. After the first sonication cell lysate was stirred and followed by another sonication of 8 min. The soluble fraction of the cell lysate was obtained by centrifuging for 30 min at 36,250 *g* at 4 °C. The supernatant was transferred to an ÄKTAxpress to carry out immobilised metal affinity chromatography (IMAC) in tandem with gel filtration. IMAC was carried out using 5 mL HisTrapTM HP NTA columns (GE Healthcare). After washing with 100 mL of washing buffer (50 mM Tris-HCl pH 8.0, 50 mM glycine, 5% glycerol, 500 mM NaCl and 20 mM imidazole), proteins were then eluted with 25 mL of elution buffer (50 mM Tris- HCl pH 8.0, 50 mM glycine, 500 mM NaCl, 500 mM imidazole, 5% (v/v) glycerol). This elution was then loaded onto a gel filtration SuperdexTM 200 HiLoadTM 26/600 column (GE Healthcare) equilibrated with gel filtration buffer (20 mM HEPES pH 7.5 and 150 mM NaCl). The gel filtration buffer for 6xHisGST-Pmk1 proteins was supplemented with 1 mM TCEP. Protein samples were separated by size and fractionated in 2 mL fractions that were analysed by SDS-PAGE to assess the presence of proteins. Fractions containing the proteins of interest where pooled and concentrated to 1 mL using VivaSpin^®^ concentrators (Sartorius) with an appropriate molecular weight cut-off. Recombinant proteins were aliquoted and frozen in liquid nitrogen for storage at -80 °C. Heterologous production and purification of MPK6 and MEK2^DD^ was performed, as previously described ^88^.

### Co-immunoprecipitation (Co-IP)

Co-IP experiments were performed to validate Pmk1 - Vts1 interactions during appressorium development. *M. oryzae* appressorium samples of 4 h development transformed were generated for *ToxA*:GFP and *VTS1-GFP*. Total protein was extracted from each frozen sample using mortar and pestle to ground into fine powder. Appressorium powder was mixed with 2x w/v ice-cold extraction buffer (10% glycerol, 25 mM Tris pH 7.5, 1 mM EDTA, 150 mM NaCl, 2% w/v PVPP, 10 mM DTT, 1x protease inhibitor cocktail (Sigma), 0.1% Tween 20 (Sigma)) and vortexed vigorously. After centrifugation at 4,200 x *g*/4 °C for 20-30 min, the supernatant was used to determine the protein concentration by the Bradford assay. The presence of each protein in the input was determined by SDS-PAGE/Western blot. *ToxA*:GFP and VTS1-GFP proteins were detected by probing the membrane with anti-GFP horseradish peroxidase (HRP)- conjugated antibody (Santa Cruz Biotechnology, USA), Pmk1 with a Phospho-p44/42 MAPK (Erk1/2) (Thr202/Tyr204) (D13.14.4E) antibody (Santa Cruz Biotechnology, Santa Cruz, CA, U.S.A.) and a HRP-conjugated anti-rabbit antibody (Abcam, UK). *M. oryzae* actin protein was used as loading control and detected with an anti-actin primary antibody (Agrisera com, Sweden) and the anti-rabbit HRP conjugated antibody.

For immunoprecipitation, 1 ug of total protein was incubated with 30 μL of GFP beads (ChromoTek, Germany) in a rotatory mixer at 4 °C. After 3 h, the beads were pelleted (800 x *g*, 1 min) and the supernatant removed. The pellet was washed and resuspended in 1 mL of IP buffer (10% glycerol, 25 mM Tris pH 7.5, 1 mM EDTA, 150 mM NaCl, 0.1% Tween 20 (Sigma)) and pelleted again by centrifugation as before. Washing steps were repeated five times. Finally, 30 μL of 1:1 dilution of SDS buffer and water supplemented with 100 mM DTT was added to the beads and incubated for 10 min at 70 °C. The beads were pelleted again, and the supernatant loaded onto SDS-PAGE gels prior to Western blotting. Membranes were probed with anti-GFP and a Phospho-p44/42 MAPK (Erk1/2) (Thr202/Tyr204) (D13.14.4E) antibody as described before. Blots membrane imaging was carried out with an ImageQuant LAS 4000 luminescent imager (GE Healthcare Life Sciences, Piscataway, NJ, U.S.A.).

### *In vitro* phosphorylation assay

For *in vitro* phosphorylation assays, 6xHis-GST tagged Pmk1 (250ng) was activated by incubation with recombinant MEK2^DD^ (250ng). Recombinant 6xHis tagged Vts1 (500ng) (500ng) was incubated with active Pmk1 in kinase buffer (25mM Tris pH 7.5, 10mM MnCl2, 1mM EGTA and 1mM DTT) in the presence of 1 mM ATP at 30 °C for 30 min. Proteins were separated by SDS-PAGE and transferred to a polyvinylidene difluoride (PVDF) membrane using a Trans-Blot turbo transfer system (Bio-Rad). PVDF membrane was blocked with 2% bovine serum albumin (BSA) in Tris-buffered saline and 1% Tween 20. His tag detection was carried using polyclonal anti-6xHis horseradish peroxidase (HRP) -conjugated antibody (Abcam). Pmk1 activated was detected using Phospho-p44/42 MAPK (Erk1/2) (Thr202/Tyr204) (Santa Cruz Biotechnology) and anti-rabbit HRP-conjugated antibodies. Pierce ECL Western Blotting Substrate (Thermo Fisher Scientific) was used for detection. Membranes were imaged using ImageQuant LAS 4000 luminescent imager (GE Life Sciences). Phosphorylation assays were analysed by mass spectrometry.

### Functional categorization of Pmk1 targets

To further understand the putative direct Pmk1 targets obtained from the MS approach, selected phosphoproteins containing a MAPK phosphorylation motif (Pxx[S/T] P or [S/T] P) were categorized based on functional annotations from Blast2GO ^89^ and Pfam ^90^.

### Protein extraction and phosphopeptide enrichment

Spores and appressoria samples were lyophilized and resuspended in extraction buffer (Urea 8M, NaCl 150 mM, Tris pH 8 100 mM, EDTA 5 Mm, aprotinin 1 µg/mL, leupeptin 2 µg/mL) for mechanical disruption using GenoGrinder 2010 (Thermo Scientific) in cold conditions (1 min at 1200 rpm). The homogenate was centrifuged for 10 min at 16,000 × *g* (Eppendorf 5415D microcentrifuge). The supernatant was used for phosphopeptide enrichment. Sample preparation started from 1.5 mg of protein extract (determined using the Bradford assay) dissolved in ammonium bicarbonate buffer containing 8 M urea. First, the protein extracts were reduced with 5 mM Tris (2-carboxyethyl) phosphine (TCEP) for 30 min at 30°C with gentle shaking, followed by alkylation of cysteine residues with 40mM iodoacetamide at room temperature for 1 hour. Subsequently, the samples were diluted to a final concentration of 1.6 M urea with 50mM ammonium bicarbonate and digested overnight with trypsin (Promega; 1:100 enzyme to substrate ratio). Peptide digests were purified using C18 SepPak columns as described before ^91^. Phosphopeptides were enriched using titanium dioxide (TiO2, GL Science) with phthalic acid as a modifier as described previously ^92^. Phosphopeptides were eluted by a pH-shift to pH 10.5 and immediately purified using C18 microspin columns (The Nest Group Inc., 5 – 60 µg loading capacity). After purification, all samples were dried in a Speedvac, stored at -80°C and re-suspended in 2% Acetonitril (AcN) with 0.1% trifluoroacetic acid (TFA) just before the mass spectrometric measurement.

### Mass-Spectrometry analysis

LC-MS/MS analysis was performed using a Orbitrap Fusion trihybrid mass spectrometer (Thermo Scientific) and a nanoflow-UHPLC system (Dionex Ultimate3000, Thermo Scientific) Peptides were trapped to a reverse phase trap column (Acclaim PepMap, C18 5 µm, 100 µm x 2 c§m, Thermo Scientific) connected to an analytical column (Acclaim PepMap 100, C18 3 µm, 75 µm x 50 cm, Thermo Scientific). Peptides were eluted in a gradient of 3-40 % acetonitrile in 0.1 % formic (solvent B) acid over 120 min followed by gradient of 40-80 % B over 6 min at a flow rate of 200 nL/min at 40°C. The mass spectrometer was operated in positive ion mode with nano-electrospray ion source with ID 0.02mm fused silica emitter (New Objective). Voltage +2200 V was applied via platinum wire held in PEEK T-shaped coupling union with transfer capillary temperature set to 275 °C. The Orbitrap, MS scan resolution of 120,000 at 400 m/z, range 300 to 1800 m/z was used, and automatic gain control (AGC) was set at 2e5 and maximum inject time to 50 ms. In the linear ion trap, MS/MS spectra were triggered with data dependent acquisition method using ‘top speed’ and ‘most intense ion’ settings. The selected precursor ions were fragmented sequentially in both the ion trap using CID and in the HCD cell. Dynamic exclusion was set to 15 sec. Charge state allowed between 2+ and 7+ charge states to be selected for MS/MS fragmentation.

### Spectral library generation and raw data processing

Peak lists in the format of Mascot generic files (.mgf files) were prepared from raw data using MSConvert package (Matrix Science). Peak lists were searched on Mascot server v.2.4.1 (Matrix Science) against either *Magnaporthe oryzae* (isolate 70-15, version 8) database, an in-house contaminants database, or *Magnaporthe oryzae* (isolate 70-15 version 8) database, Uniprot Rice database (UP000007015; *Oryza sativa subspecies indica*, strain: cv. 93-11) and an in-house contaminants database. Tryptic peptides with up to 2 possible mis-cleavages and charge states +2, +3, +4, were allowed in the search. The following modifications were included in the search: oxidized methionine, phosphorylation on Serine, Threonine, Tyrosine as variable modification and carbamidomethylated cysteine as static modification. Data were searched with a monoisotopic precursor and fragment ions mass tolerance 10ppm and 0.6 Da respectively. Mascot results were combined in Scaffold v. 4 (Proteome Software) and exported in Excel (Microsoft Office).

### Label free quantification at MS1 level

Peptide quantification was performed as described recently ^93^ with the following modifications. Raw data files were processed using Proteome Discoverer 2.5 (Thermo Fisher Scientific) and searched against an in-house constructs and contaminants database and the *Magnaporthe oryzae* (isolate 70-15 version 8) database. The processing workflow was made up of the Sequest HT search engine, Percolator (for target/decoy selection) and IMP-ptmRS (to calculate modification site probabilities). Tryptic peptides with up to 2 possible mis-cleavage and charge states +2, +3 were allowed in the search and the follow modifications were included in the search: carbamidomethylated Cysteine (fixed), oxidized Methionine (variable) and phosphorylated Serine, Threonine and Tyrosine (variable). Data were searched with a monoisotopic precursor and fragment ion mass tolerance 10 ppm and 0.6 Da respectively. Peptides were quantified using the ‘basic modification analysis’ consensus workflow provided by Proteome Discoverer 2.5 and expressed as abundance ratios. Peptides in the Peptide groups tab in the results files were filtered for ‘phospho’ and reliable and detectable ‘quan’ values. Threshold for differential phosphopeptides was set at minimum 2-fold change in abundance ratio and an adjusted abundance ratio p-value of less than 0.05. Data for Peptide groups were exported to Excel and processed in R.

### Phosphosite conservation analysis

To determine the conservation of phosphosites across species, a list of 41 fungi of various lifestyles was prepared, and the protein sequences for a given assembly downloaded from the sources listed in Supplemental Table 1. These were then used in Orthofinder version 2.3.7 ^94^. Running diamond 2.0.14 ^95^ to compute orthogroups and species trees. Each *M. oryzae* phos- site was taken in turn and the orthologues of the source protein were compared to it in turn using blastp from BLAST+2.9.0 ^96^. If any matches were found according to BLAST defaults, the best HSP (by bitscore) was retained and the HSP in the orthologue was extended to match the full range of the length of the *M. oryzae* phosphopeptide sequence. If then the phos-site lies in the range the HSP of the orthologue was then checked to see whether the corresponding residue in the orthologue has a match to the *M. oryzae* phos-site residue.

### Clustering of phosphosites by conservation

For each phos-site and query fungal species the proportion of M.oryzae sites matched was calculated and tabluted for *k*-means clustering. The value of *k* was determined by scanning values of k from 2 to 50 and calculating the variance as Within Sum of Squares at each *k*. The variance in clusters stopped decreasing noticeably at about k = 9 and that was taken as the value of *k* for final clustering. *K* means clustering was performed using the factoextra package^97^ kmeans function in R ^98^ version 4.2.0. Data preparation was performed in R using the tidyverse packages ^99^. Heatmaps were prepared using ComplexHeatmap^100^. Phylogenetic trees were analyzed in ape^101^, dendextend ^102^ and rendered in ggtree^103^.

### Clustering and Gene Ontology Analysis

*M. oryzae* Gene Ontology analysis was performed using the version MG8 annotations from ENSEMBL BioMart. Cluster enrichment computations were performed in the R package clusterProfiler 4.6.2^104^ at a p-value of <= 0.05 with Benjamini-Hochberg corrections for multiple hypothesis tests.

### Parallel Reaction Monitoring (PRM)

Peptide quantitation was performed using Parallel Reaction Monitoring (PRM) as described previously ^105^. Briefly, mass to charge ratios (m/z) corresponding to selected phospho-peptides were monitored and filtered by the first quadrupole and fragment ions were scanned out in the orbitrap mass analyzer over the duration of the elution profile. The PRM assay also included a selection of control peptides having similar relative intensities in each sample and used to measure relative phospho-peptide content (Supplementary Table 1). Raw data were peak picked and searched against the data bases on the Mascot server as described above and combined with chromatographic profiles in Skyline ^106^ to determine individual peptide intensities. Extracted phospho-peptides intensity were normalized against the summed control peptide intensities to correct for differences in phospho-peptide yield. The assay was performed once for each of three biological replicates and results were subjected to differential phosphosite analysis.

### Differential phosphosite analysis

To determine whether phosphosites were differentially abundant between samples we prepared PRM data by replacing missing values with the lowest observed intensity in that replicate and then performed a bootstrap *t*-test with 1000 bootstrap resamples with replacement for each phosphosite using the MKInfer package ^107^. We used *p* <= 0.05 as a threshold for differential abundance.

## SUPPLEMENTAL INFORMATION TITLES AND LEGENDS

**Supplemental Figure 1.**
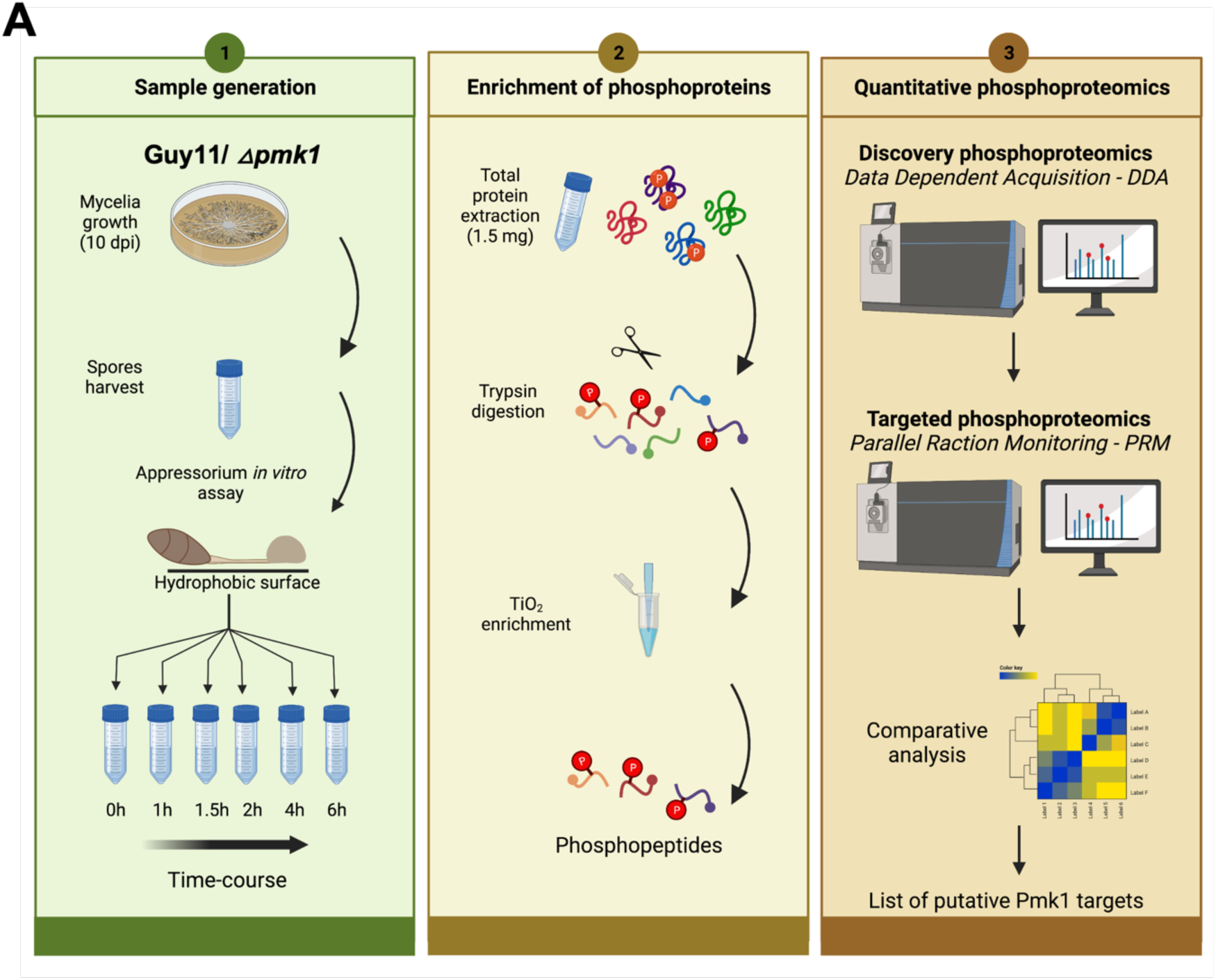
Phosphoproteomics experimental workflow and data analysis pipeline to determine the phosphorylation landscape during infection-related development in *M. oryzae*. Flowchart showing the experimental strategy during sample generation, phosphopeptide enrichment and quantitative phosphoproteomic analysis to determine the phosphorylation landscape of early infection and the Pmk1 targets.

**Supplemental Figure 2.**
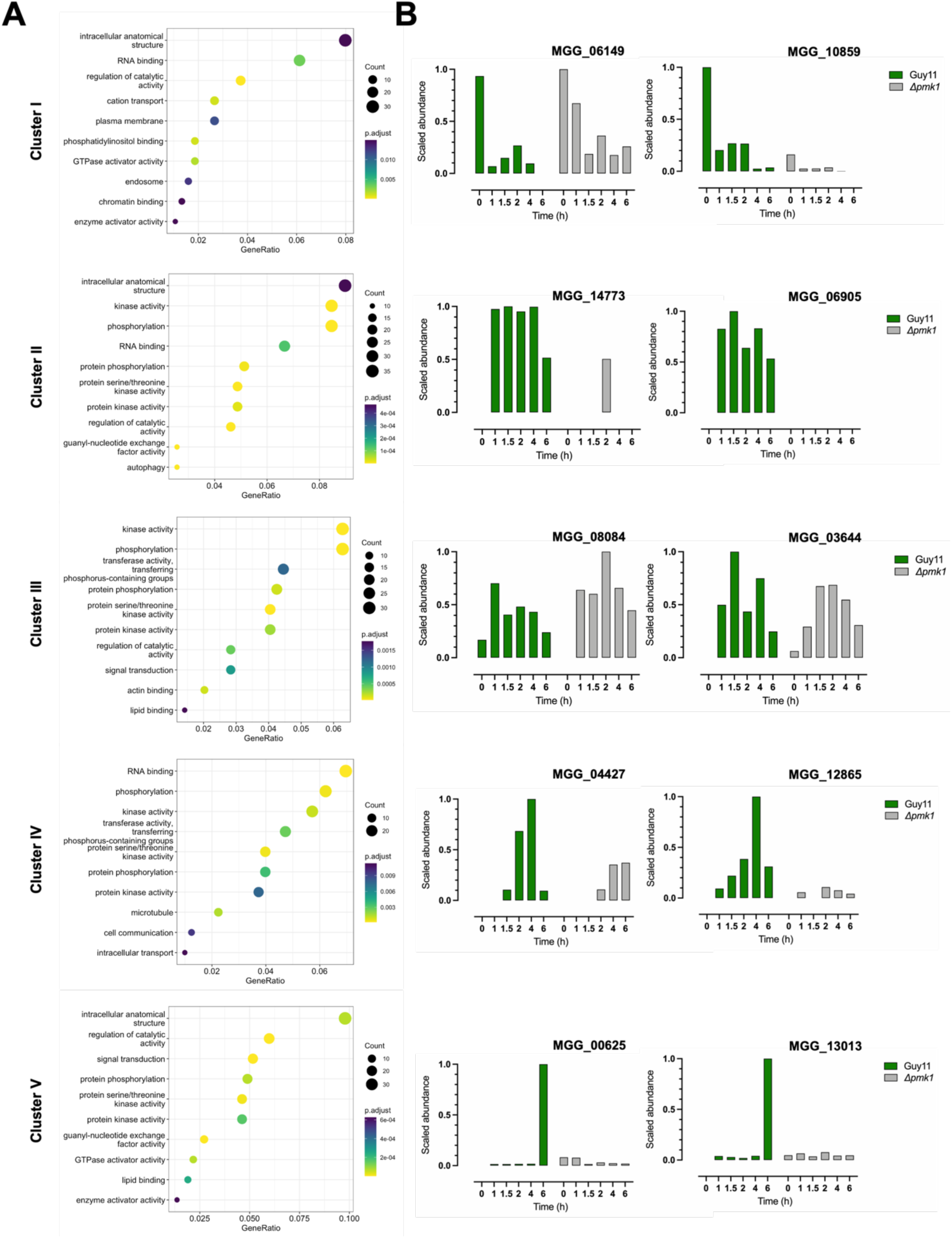
GO-term enrichment for differentially phosphorylated proteins (MS1 analysis). **(A)** GO term analysis of differentially phosphopeptides analyzed by MS1, categorized according to the time point at which they passed significance threshold. **(B)** Relative phosphorylation for representative phosphopeptides in each defined cluster. Cluster I, MGG_05433 and MGG_10859; Cluster II, MGG_14773 and MGG_06905; Cluster III, MGG_08084 and MGG_03644; Cluster IV, MGG_04427 and MGG_12865; and Cluster V, MGG_00625 and MGG_13013.

**Supplemental Figure 3.**
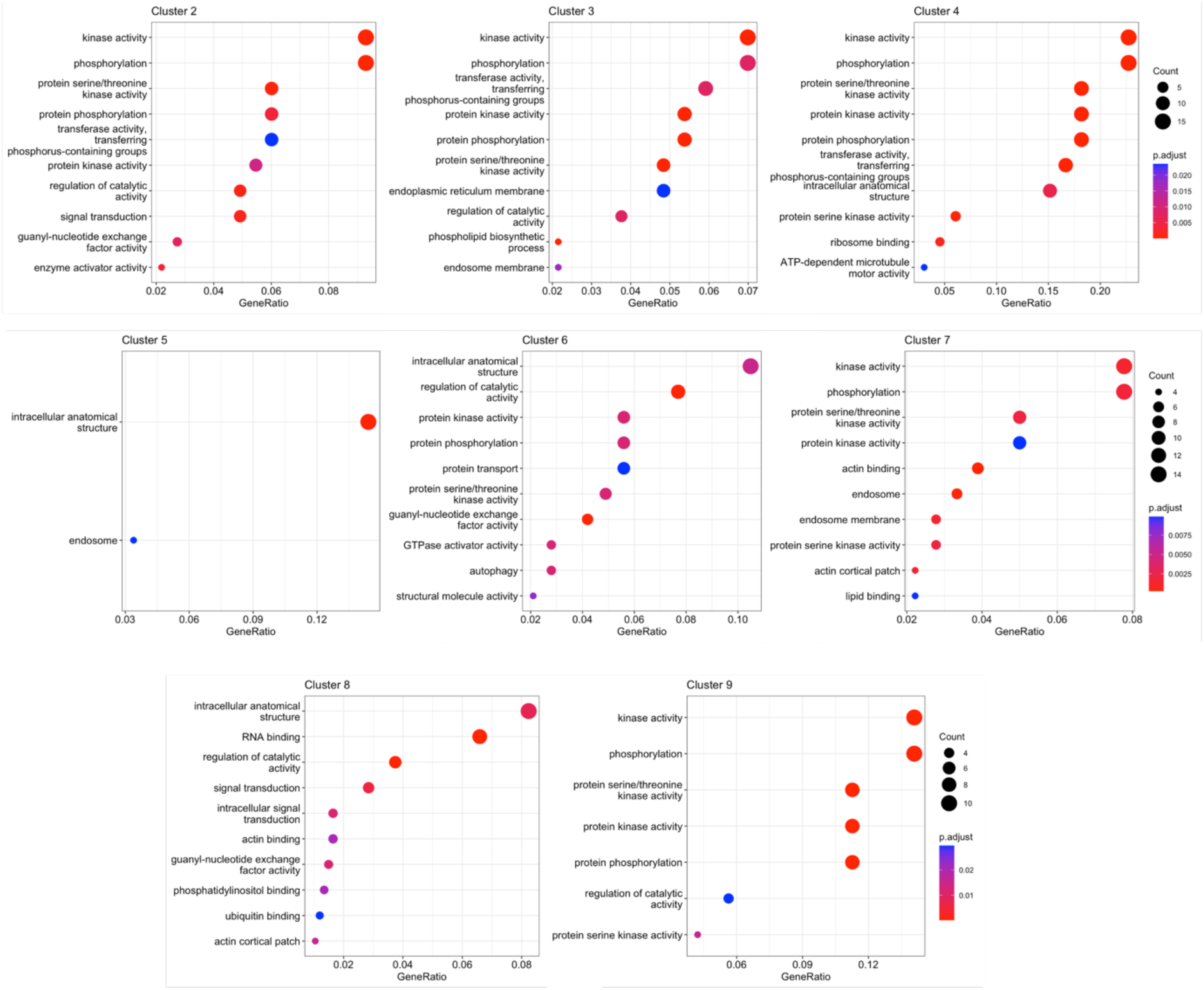
GO-term enrichment for the Conserved Phosphorylated Residue (CPR) groups. GO term analysis of evolutionary conserved phosphopeptides analyzed by MS1, categorized according to the time point at which they passed significance threshold.

**Supplementary Figure 4.**
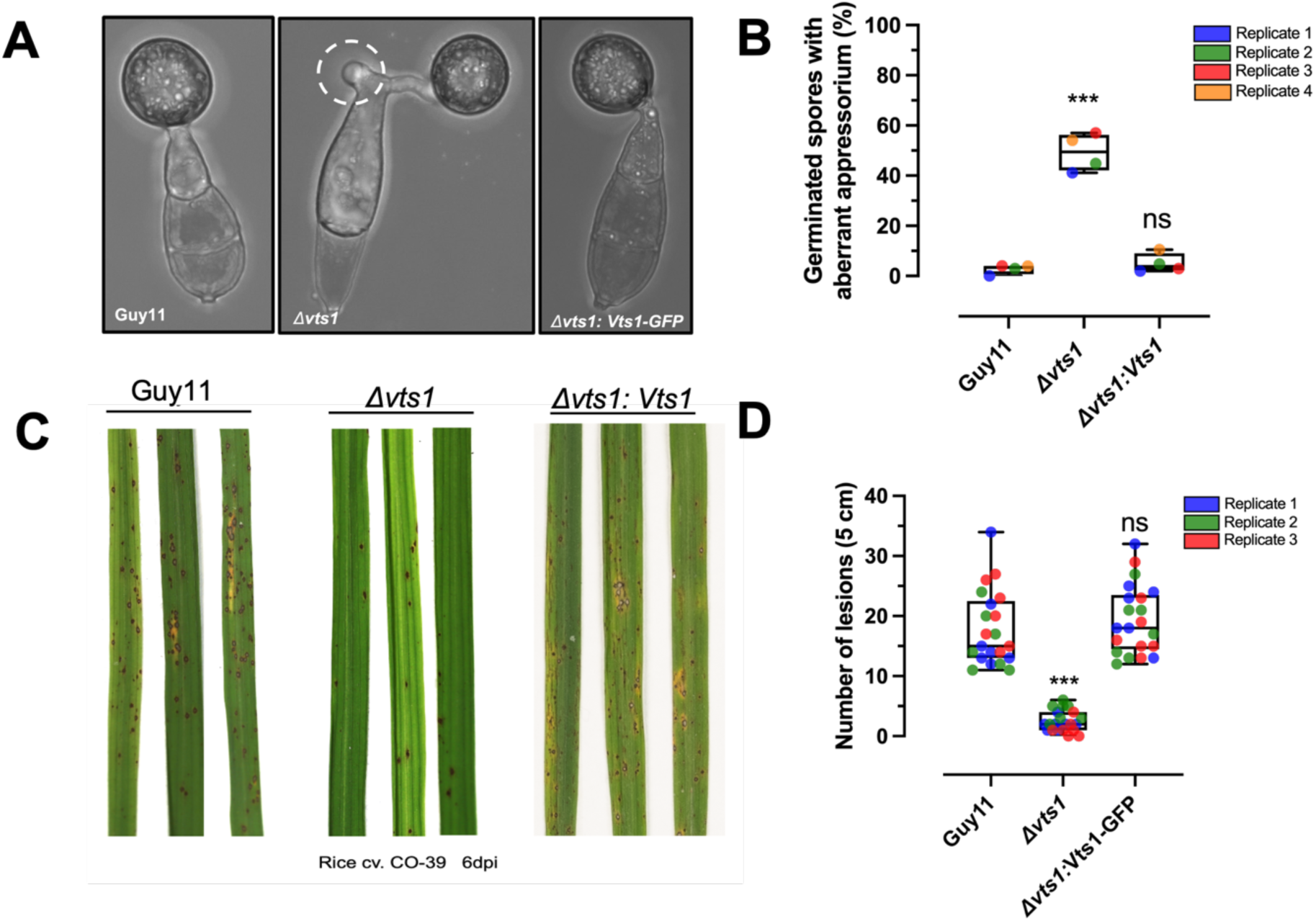
Phosphorylation profile of early appressorium development pathways. Heat map to show relative intensities of differentially phosphorylated peptides in Guy11 and 11*pmk1* clustered by signalling pathway. MGG number, gene name and phosphosite detected are shown.

**Supplementary Figure 5.**
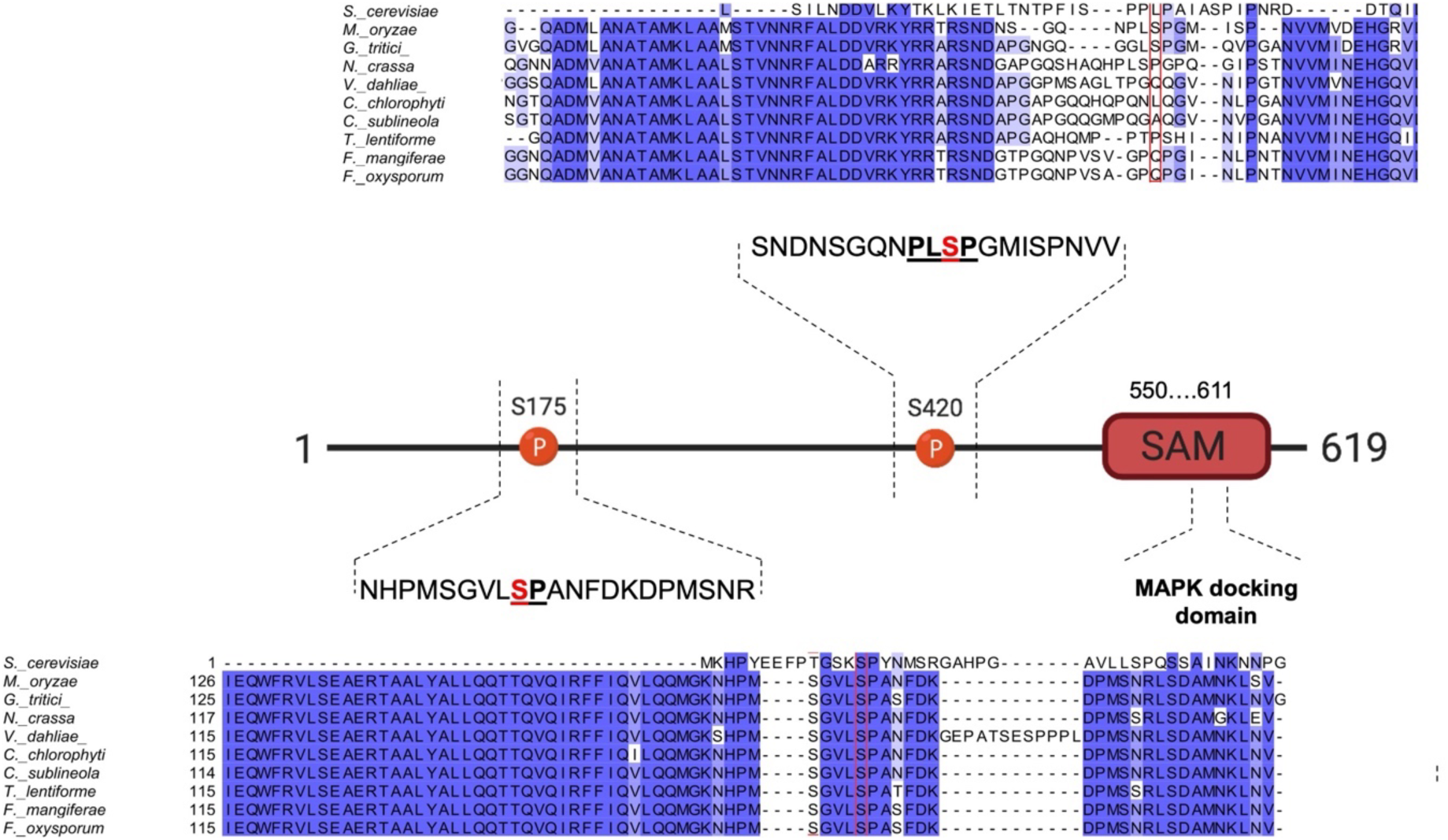
Quantitative phosphoproteomics defines putative targets of the Pmk1 MAPK pathway. Heat map to show relative intensities of 181 differentially phosphorylated peptides in Guy11 and 11*pmk1.* MGG number, gene name and phosphosite detected are shown.

**Supplemental Figure 6.**
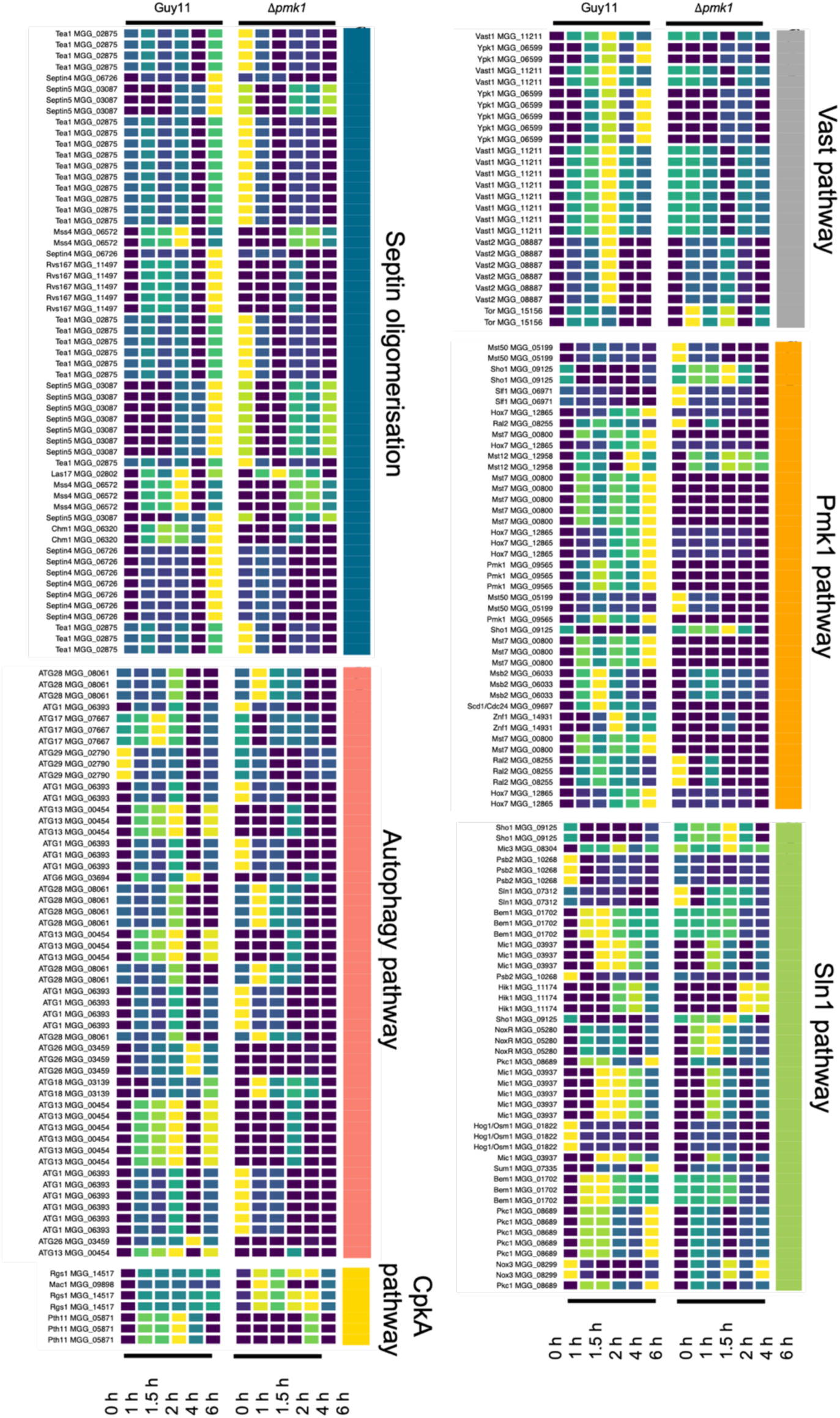
Experimental design for parallel reaction monitoring of the Pmk1 analogue-sensitive mutant of *M. oryzae* during the early stages of appressorium development. The *pmk1^AS^* conditional mutant was incubated on a hydrophobic surface for 1, 2,3 and 4 h in the presence or absence of the ATP analogue 1NaPP1.

**Supplemental Figure 7.**
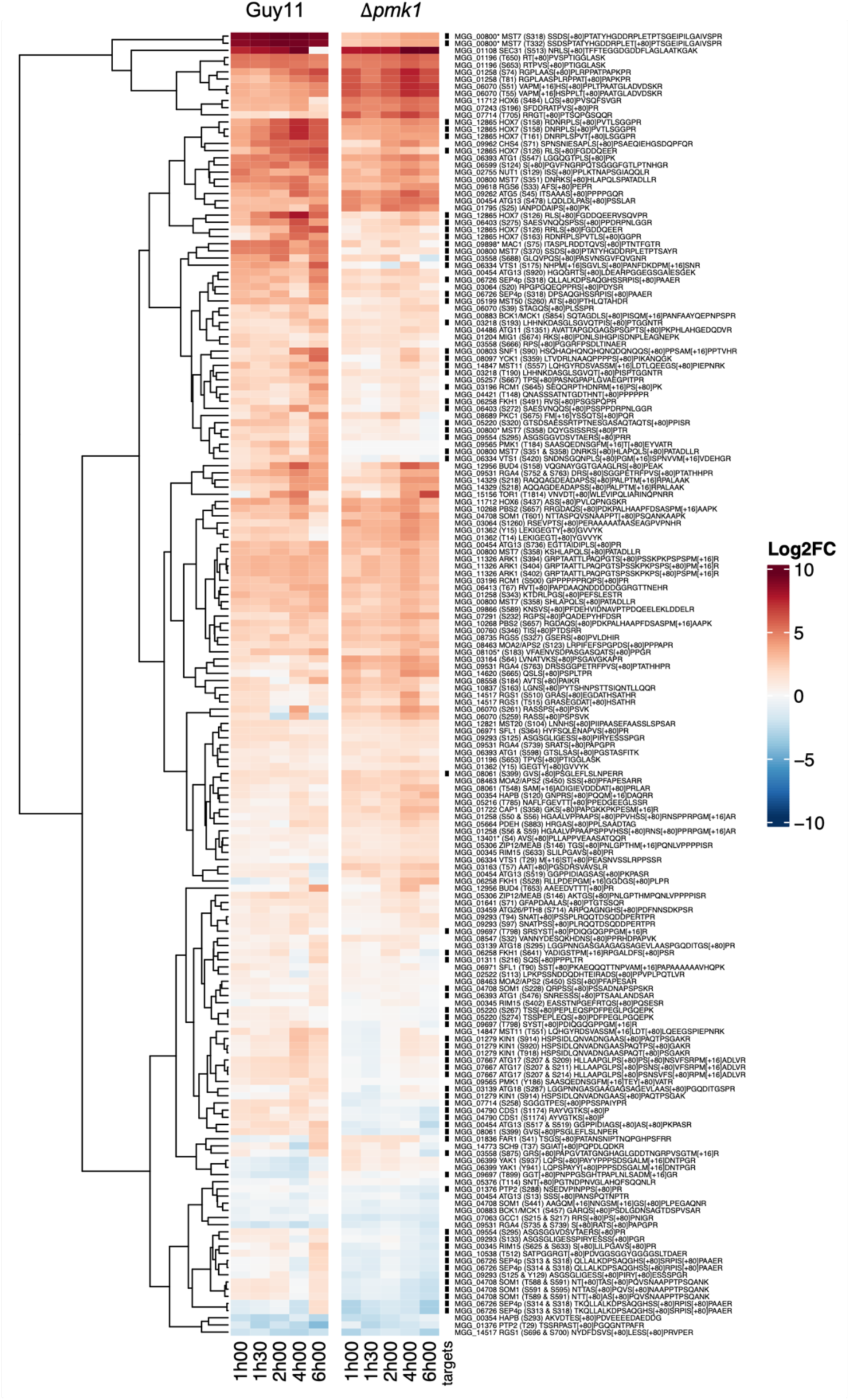
MEK2DD activates *in vitro* Pmk1 on its TEY motif. **(A)** Western blot analysis of *in vitro* phosphorylation experiment between MEK2^DD^ (N-terminally tagged with 6xHis) and Pmk1 (N-terminally tagged with GST). The previously reported MEK2^DD^ phosphorylation of MPK6 (N-terminally tagged with 6xHis) was used as a positive control. Proteins were immunoblotted with appropriate antisera (listed on the right). Arrows indicate expected band sizes. **(B)** Phosphopeptides identified by LC-MS for the *in vitro* kinase assay. **(C)** Phosphorylation sites (in red) identified by LC-MS on the Pmk1 MAPK. **(D)** Phosphopeptides identified by LC- MS for the *in vitro* kinase assay. **(E)** Phosphorylation sites (in red) identified by LC-MS on Vts1.

**Supplemental Figure 8.**
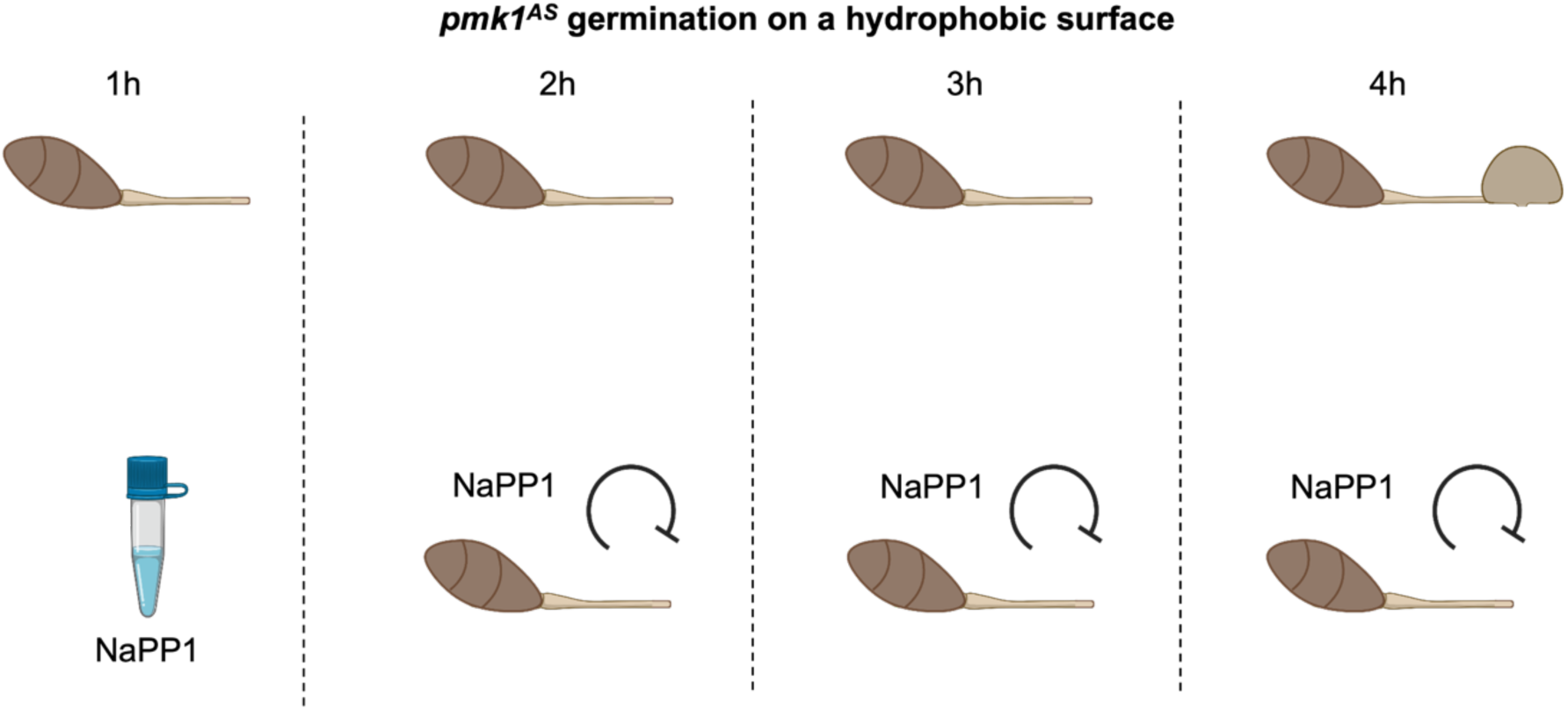
Southern blot and whole genome sequencing confirm VTS1 null mutant. **(A)** A probe of 1046 bp was generated to hybridise to the *VTS1* gene in its 5’ UTR region. **(B)** Genomic DNA of the putative transformants was digested with *Hin*dIII, gel fractionated, and transferred to Hybond-NX. **(C)** Southern blot analysis showing a single band of 4468 bp for positive null mutants (KO) and 1425 bp for wild-type strains (WT). The blot was probed with 1046 bp DNA fragment specific to *VTS1*. **(D)** Bioinformatic analysis of one positive 11*vts1* null mutant showing the absence of coverage (reads) for the *VTS1* gene due to the presence of the *HPH* cassette inserted by the split marker strategy.

**Supplemental Figure 9.**
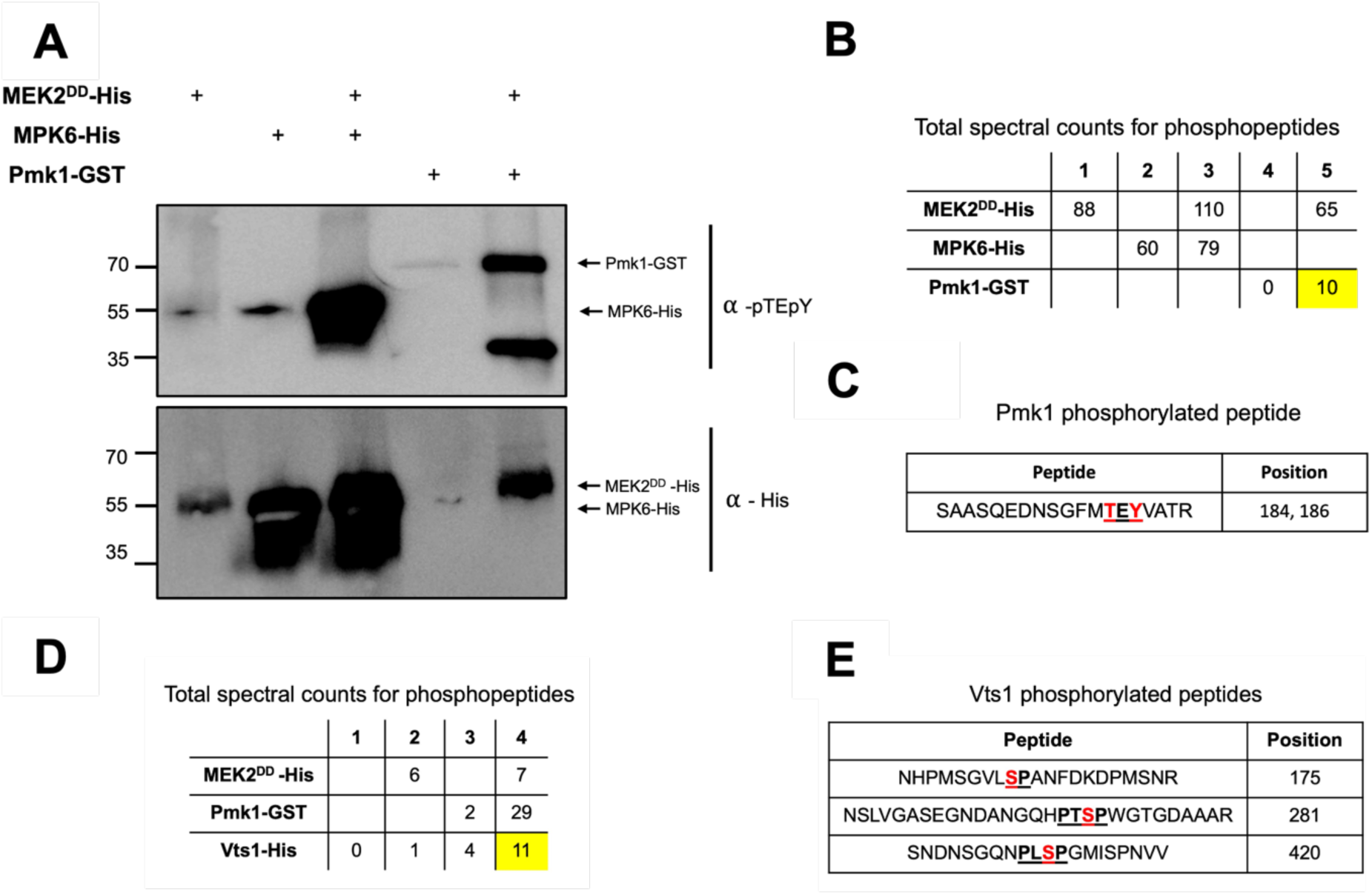
A Pmk1-dependent Vts1 phosphorylation is necessary for rice blast disease. **(A)** Micrographs to show appressorium development of Guy11, Δ*vts1 and* 11*vts1:*VTS1-GFP strains. Conidia were harvested from Guy11 and Δ*vts1* mutants, inoculated onto glass coverslips, and observed at 24h. Scale bar= 10μm. **(B)** Bar chart showing the frequency of conidial germination from one and two cells. Three biological replicates were carried out with 100 appressoria recorded per replicate. \**P* < 0.05: \*\**P* < 0.01; \*\*\**P* < 0.001 represent significant differences using an unpaired two-tailed Student’s *t*-test. Data are from four biological replicates. **(C)** Two-week-old seedlings of rice cultivar CO-39 were inoculated with equal amounts of conidial suspensions of Guy11 and Δ*vts1* containing 10^5^ conidia mL^-1^ in 0.2% gelatine. Seedlings were incubated for 6 days to develop blast disease at 26 °C and 90 % humidity. **(D)** Scatter chart to show the number of disease lesions in Guy11 and two independent 11*vts1* mutants. Horizontal line represents the mean, and the error bar is the standard deviation Data points are shown from three biological replicates in different colours (red, blue, green).

**Supplemental Figure 10.**
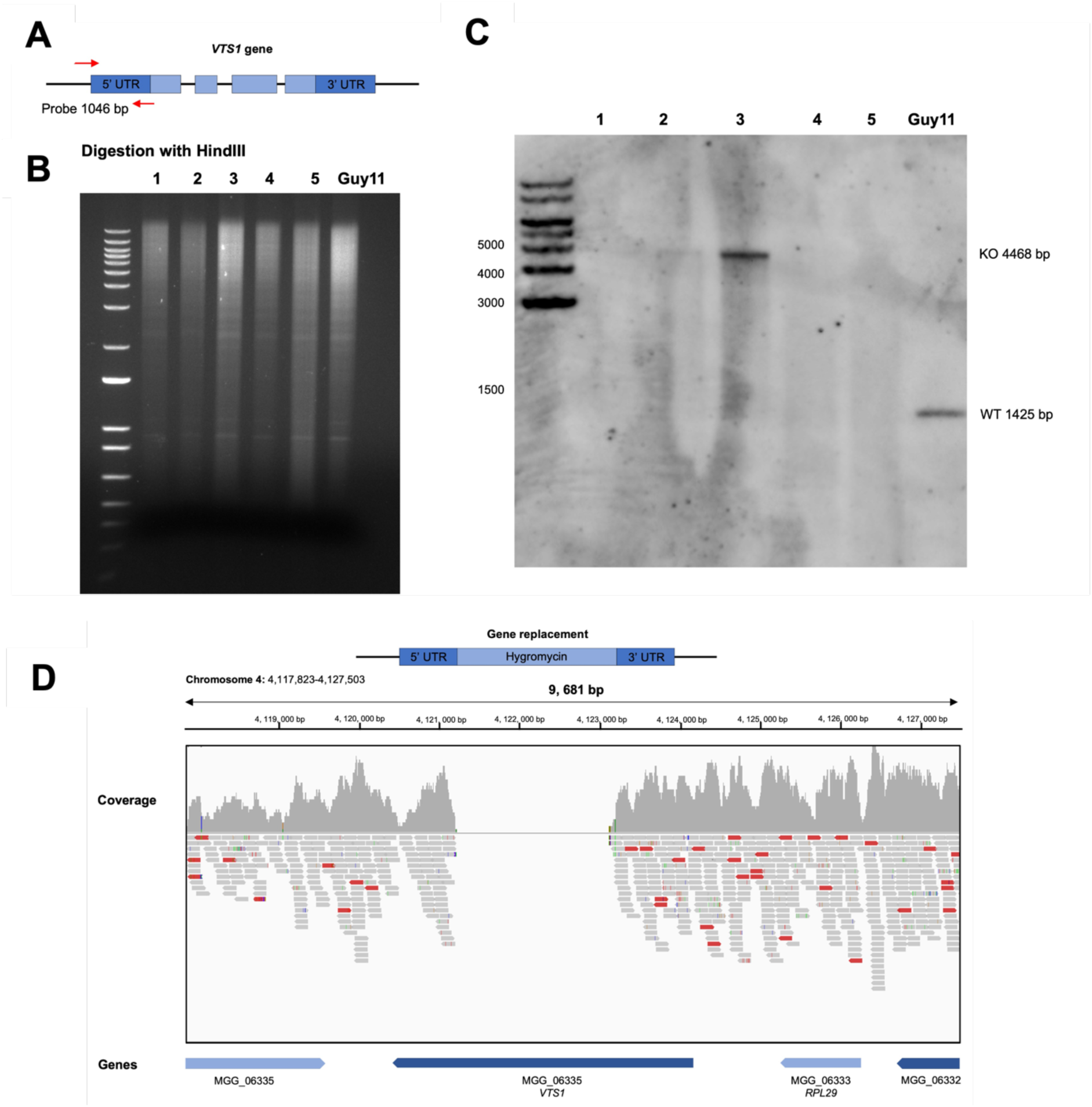
The Vts1 S175 residue is present in a conserved region among filamentous fungi. Schematic representation to show arrangement of each phosphorylation site identified for Vts1 and its conservation. Alignments of neighbouring regions surrounding Vts1 S175 and S420 from different filamentous fungi were carried out using ClustalX.

**Supplementary Table 1.** Name, assembly version and source of proteome sequences used for phosphorylation site conservation analysis.

| Name | Version | Source |
| --- | --- | --- |
| *M. oryzae* | MG8 | https://ftp.ebi.ac.uk/ensemblgenomes/pub/release-56/fungi/fasta/magnaporthe_oryzae/pep/Magnaporthe_oryzae.MG8.pep.all.fa.gz |
| *A. alternata* | Altal1 | https://ftp.ensemblgenomes.ebi.ac.uk/pub/fungi/release-56/fasta/fungi_ascomycota3_collection/alternaria_alternata_gca_001642055/pep/Alternaria_alternata_gca_001642055.Altal1.pep.all.fa.gz |
| *C. fruticola* | ASM977102v1 | https://ftp.ensemblgenomes.ebi.ac.uk/pub/fungi/release-56/fasta/fungi_ascomycota4_collection/colletotrichum_fruticola_gca_009771025/pep/Colletotrichum_fruticola_gca_009771025.ASM977102v1.pep.all.f a.gz |
| *C. gloeosporioides* | GCA_000319635.1 | https://ftp.ensemblgenomes.ebi.ac.uk/pub/fungi/release-56/fasta/colletotrichum_gloeosporioides/pep/Colletotrichum_gloeosporioides.GCA_000319635.1.pep.all.fa.gz |
| *C. higginsianum* | GCA_000313795.2 | https://ftp.ensemblgenomes.ebi.ac.uk/pub/fungi/release-56/fasta/colletotrichum_higginsianum/pep/Colletotrichum_higginsianum.GCA_000313795.2.pep.all.fa.gz |
| *B. graminis* | EF2 | https://ftp.ensemblgenomes.ebi.ac.uk/pub/fungi/release-56/fasta/blumeria_graminis/pep/Blumeria_graminis.EF2.pep.all.fa.gz |
| *P. graminis* | ASM14992v1 | https://ftp.ensemblgenomes.ebi.ac.uk/pub/fungi/release-56/fasta/puccinia_graminis/pep/Puccinia_graminis.ASM14992v1.pep.all.fa.gz |
| *U. maydis* | Umaydis521_2.0 | https://ftp.ensemblgenomes.ebi.ac.uk/pub/fungi/release-56/fasta/ustilago_maydis/pep/Ustilago_maydis.Umaydis521_2.0.pep.all.fa.gz |
| *P. pachyrhizi* | PpacPPUFV02 | https://genome.jgi.doe.gov/portal/PpacPPUFV02/download/PpacPPUFV02_GeneCatalog_proteins_20210128.aa.fasta.gz |
| *B. oryzae* | Cochliobolus_miyabeanus_v1 | https://ftp.ensemblgenomes.ebi.ac.uk/pub/fungi/release-56/fasta/fungi_ascomycota1_collection/bipolaris_oryzae_atcc_44560_gca_000523455/pep/Bipolaris_oryzae_atcc_44560_gca_000523455.Cochliobolus_miyabeanus_v1.0.pep.all.fa.gz |
| *B. sorokiniana* | Cocsa1 | https://ftp.ensemblgenomes.ebi.ac.uk/pub/fungi/release-56/fasta/fungi_ascomycota1_collection/bipolaris_sorokiniana_nd90pr_gca_000338995/pep/Bipolaris_sorokiniana_nd90pr_gca_000338995.Cocsa1.pep.all.f a.gz |
| *B. cinerea* | ASM83294v1 | https://ftp.ensemblgenomes.ebi.ac.uk/pub/fungi/release-56/fasta/botrytis_cinerea/pep/Botrytis_cinerea.ASM83294v1.pep.all.fa.gz |
| *C. heterostrophus* | CocheC5_3 | https://genome.jgi.doe.gov/portal/CocheC5_3/download/CocheC5_1_GeneModels_FilteredModels1_aa.fasta.gz |
| *P. striiformis* | PST-130_1.0 | https://ftp.ensemblgenomes.ebi.ac.uk/pub/fungi/release-56/fasta/puccinia_striiformis/pep/Puccinia_striiformis.PST-130_1.0.pep.all.fa.gz |
| *S. cerevisiae* | ScR64-1-1 | https://ftp.ebi.ac.uk/ensemblgenomes/pub/release-56/fungi/fasta/saccharomyces_cerevisiae/pep/Saccharomyces_cerevisiae.R64-1-1.pep.all.fa.gz |
| *F. oxysporum-5176* | GCA_000222805.1 | https://ftp.uniprot.org/pub/databases/uniprot/current_release/knowledgebase/reference_proteomes/Eukaryota/UP000002489/UP000002489_660025.fasta.gz |
| *A. nidulans* | ASM1142v1 | https://ftp.ensemblgenomes.ebi.ac.uk/pub/fungi/release-56/fasta/aspergillus_nidulans/pep/Aspergillus_nidulans.ASM1142v1.pep.all.f a.gz |
| *S. pombe* | ASM294v2 | https://ftp.ensemblgenomes.ebi.ac.uk/pub/fungi/release-56/fasta/schizosaccharomyces_pombe/pep/Schizosaccharomyces_pombe.ASM294v2.pep.all.fa.gz |
| *N. crassa* | NC12 | https://ftp.ensemblgenomes.ebi.ac.uk/pub/fungi/release-56/fasta/neurospora_crassa/pep/Neurospora_crassa.NC12.pep.all.fa.gz |
| *C. albicans* | SC5314 v22 | http://candidagenome.org/download/sequence/C_albicans_SC5314/Assembly22/current/C_albicans_SC5314_version_A22-s07-m01-r182_default_protein.fasta.gz |
| *A. fumigatus* | ASM265v1 | https://ftp.ensemblgenomes.ebi.ac.uk/pub/fungi/release-56/fasta/aspergillus_fumigatus/pep/Aspergillus_fumigatus.ASM265v1.pep.all.fa.gz |
| *C. neoformans* | ASM9104v1 | https://ftp.ensemblgenomes.ebi.ac.uk/pub/fungi/release-56/fasta/cryptococcus_neoformans/pep/Cryptococcus_neoformans.ASM9104v1.pep.all.fa.gz |
| *H. capsulatum* | GCA000170615v1 | https://ftp.ensemblgenomes.ebi.ac.uk/pub/fungi/release-56/fasta/histoplasma_capsulatum/pep/Histoplasma_capsulatum.GCA000170615v1.pep.all.fa.gz |
| *F. oxysporum-2* | FO2 | https://ftp.ensemblgenomes.ebi.ac.uk/pub/fungi/release-56/fasta/fusarium_oxysporum/pep/Fusarium_oxysporum.FO2.pep.all.fa.gz |
| *C. purpurea* | ASM34735v1 | https://ftp.ensemblgenomes.ebi.ac.uk/pub/fungi/release-56/fasta/fungi_ascomycota1_collection/claviceps_purpurea_20_1_gca_000347355/pep/Claviceps_purpurea_20_1_gca_000347355.ASM34735v1.pep.all.fa.gz |
| *A. brassicicola* | Albra1 | https://genome.jgi.doe.gov/portal/Altbr1/download/Alternaria_brassicicola_proteins.fasta.gz |
| *A. flavus* | Aspfl2_3 | https://genome.jgi.doe.gov/portal/Aspfl2_3/download/Aspfl2_3_GeneCatalog_proteins_20200207.aa.fasta.gz |
| *C. chrysosperma* | Cytch1 | https://genome.jgi.doe.gov/portal/Cytch1/download/Cytch1_GeneCatalog_proteins_20171014.aa.fasta.gz |
| *F. graminearum* | RR1 | https://ftp.ensemblgenomes.ebi.ac.uk/pub/fungi/release-56/fasta/fusarium_graminearum/pep/Fusarium_graminearum.RR1.pep.all.fa.gz |
| *F. verticillioides* | ASM331697v2 | https://ftp.ensemblgenomes.ebi.ac.uk/pub/fungi/release-56/fasta/fungi_ascomycota5_collection/fusarium_verticillioides_gca_003316975/pep/Fusarium_verticillioides_gca_003316975.ASM331697v2.pep.all.fa.gz |
| *Z. tritici* | MG2 | https://ftp.ensemblgenomes.ebi.ac.uk/pub/fungi/release-56/fasta/zymoseptoria_tritici/pep/Zymoseptoria_tritici.MG2.pep.all.fa.gz |
| *P. oxalicum* | pde_v1.0 | https://ftp.ensemblgenomes.ebi.ac.uk/pub/fungi/release-56/fasta/fungi_ascomycota1_collection/penicillium_oxalicum_114_2_gca_000346795/pep/Penicillium_oxalicum_114_2_gca_000346795.pde_v1.0.pep.all.fa.gz |
| *P. teres* | ASM972866v1 | https://ftp.ensemblgenomes.ebi.ac.uk/pub/fungi/release-56/fasta/fungi_ascomycota4_collection/pyrenophora_teres_f_teres_gca_0097 |
|  |  | 28665/pep/Pyrenophora_teres_f_teres_gca_009728665.ASM972866v1.pep.all.fa.gz |
| *S. sclerotiorum* | ASM185786v1 | https://ftp.ensemblgenomes.ebi.ac.uk/pub/fungi/release-56/fasta/fungi_ascomycota3_collection/sclerotinia_sclerotiorum_1980_uf_70_gca_001857865/pep/Sclerotinia_sclerotiorum_1980_uf_70_gca_001857865.ASM185786v1.pep.all.fa.gz |
| *S. turcica* | Settu3 | https://genome.jgi.doe.gov/portal/Settu3/download/Settu3_GeneCatalog_proteins_20170818.aa.fasta.gz |
| *S. nodorum* | Stano2 | https://genome.jgi.doe.gov/portal/Stano2/download/Stano2_GeneCatalog_proteins_20110506.aa.fasta.gz |
| *U. virens* | ASM96522v2 | https://ftp.ensemblgenomes.ebi.ac.uk/pub/fungi/release-56/fasta/fungi_ascomycota3_collection/ustilaginoidea_virens_gca_000965225/pep/Ustilaginoidea_virens_gca_000965225.ASM96522v2.pep.all.fa.gz |
| *V. mali* | ASM81815v1 | https://ftp.ensemblgenomes.ebi.ac.uk/pub/fungi/release-56/fasta/fungi_ascomycota3_collection/valsa_mali_gca_000818155/pep/Valsa_mali_gca_000818155.ASM81815v1.pep.all.fa.gz |
| *V. dahliae* | VDAG_JR2v.4.0 | https://ftp.ensemblgenomes.ebi.ac.uk/pub/fungi/release-56/fasta/verticillium_dahliaejr2/pep/Verticillium_dahliaejr2.VDAG_JR2v.4.0.pep.all.fa.gz |
| *P. indica* | Pirin1 | https://genome.jgi.doe.gov/portal/Pirin1/download/Pirin1_GeneCatalog_proteins_20111203.aa.fasta.gz |
| *R. irregularis* | ASM43914v3 | https://ftp.ensemblgenomes.ebi.ac.uk/pub/fungi/release-56/fasta/fungi_mucoromycota1_collection/rhizophagus_irregularis_daom_181602_gca_000439145/pep/Rhizophagus_irregularis_daom_181602_gca_000439145.ASM43914v3.pep.all.fa.gz |
| *F. oxysporum-C.alt* | FCALT_unknown | Berasategui et al., Curr Biology 2022 [10.1016/j.cub.2022.07.065](https://doi.org/10.1016/j.cub.2022.07.065) |

**Supplementary Table 2:**
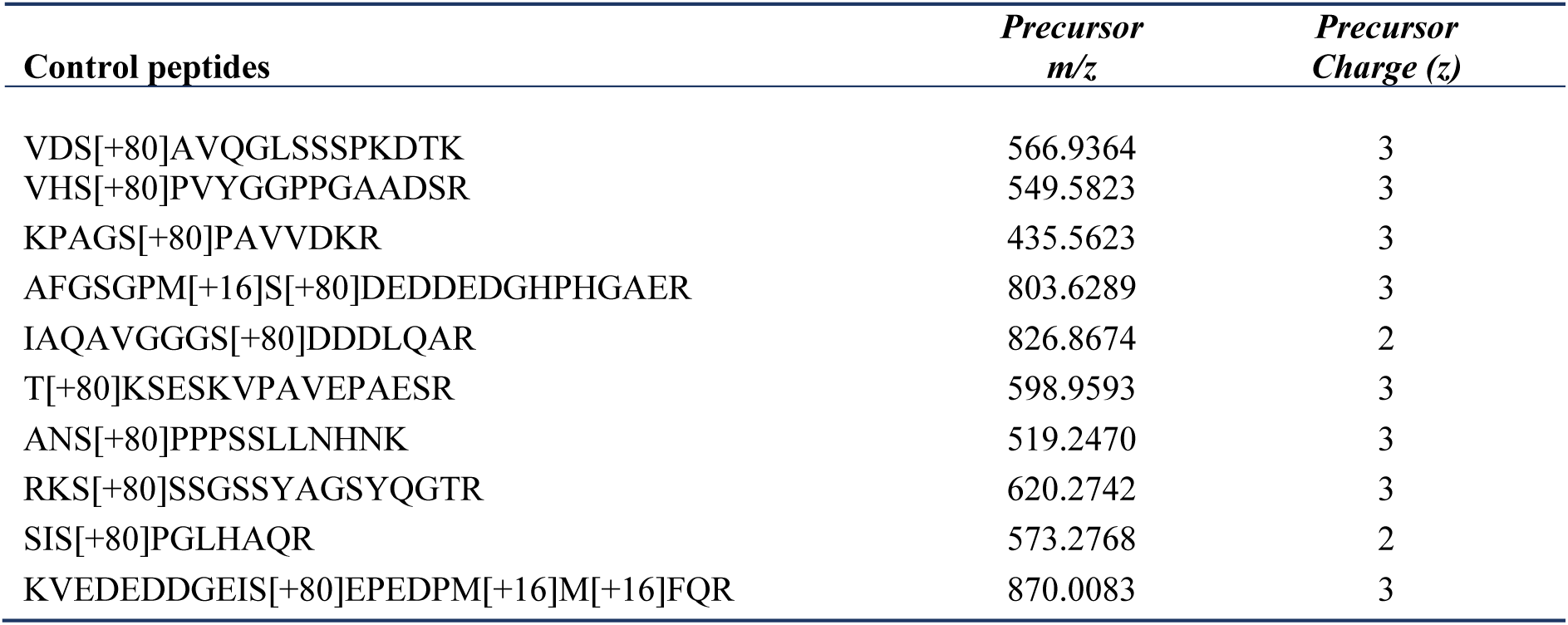
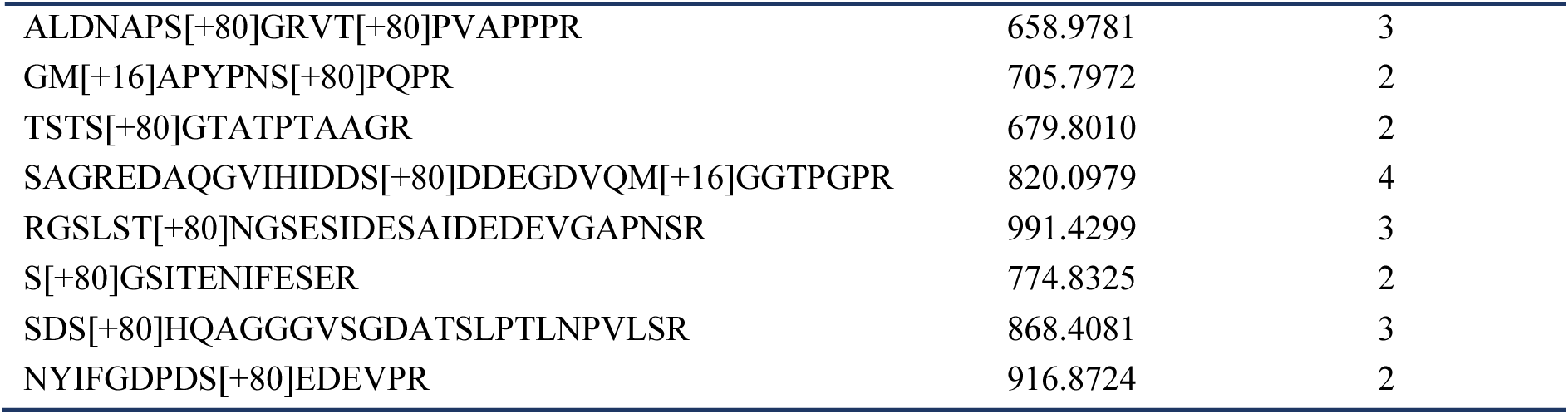
Control peptides used in PRM assay.

